# A Pan-Cancer Single-Cell Atlas to Evaluate Tumor Identity, Cell Line Concordance, and Dependency Mapping

**DOI:** 10.64898/2026.02.14.705396

**Authors:** Rosyli F Reveron-Thornton, James P Agolia, Chuner Guo, Maria Korah, Chia-Hsin Hsu, Peter Y Xie, Renceh AB Flojo, Andrea E Delitto, Amanda Gonçalves, Angela D Tabora, Michael Januszyk, Victoria E Sanchez, Kevin Nee, Biren Reddy, Wesley Bobst, Byrne Lee, George A Poultsides, Amanda R Kirane, Derrick C Wan, Jeffrey A Norton, Edgar G Engleman, Aaron M Newman, Michael T Longaker, Deshka S Foster, Daniel Delitto

## Abstract

Bulk RNA sequencing enables pan-cancer transcriptional analyses, but obscures cancer cell-specific programs due to admixture with nonmalignant cells, thereby limiting direct comparison between experimental models and primary tumors. Single-cell RNA sequencing (scRNA-seq) overcomes these limitations; however, the biological interpretability of public datasets is often compromised by variable data quality, inconsistent annotation, and atlas-scale aggregation strategies that prioritize data volume over biological coherence. We therefore developed a stringent integration framework that prioritizes representative malignant transcriptional states. Using Mahalanobis distance-based selection within batch-corrected latent space, we constructed a pan-cancer atlas comprising 135,424 high-quality malignant cells from 499 samples across 36 adult and pediatric cancers. Atlas-derived cancer signatures were used to determine tumor–cell line concordance and project ElasticNet models trained on DepMap CRISPR screens to infer cancer-specific gene dependencies. The scTumor Atlas establishes a scalable framework for tumor identity inference, cancer cell line benchmarking, and systematic identification of genetic vulnerabilities.

## Introduction

Bulk RNA sequencing (RNA-seq) has enabled expansive, pan-cancer transcriptional analyses and remains foundational for large cohort studies. However, bulk profiles represent an average signal across heterogeneous tumor ecosystems, where malignant cells coexist with stromal and immune populations. This cellular admixture obscures cancer cell-specific transcriptional programs and complicates direct comparison between human tumors and experimental models. Numerous informatic strategies have been proposed to mitigate this limitation, including deconvolution^1,2^ and inferred tumor purity correction.^3,4^ While valuable, these approaches rely on assumptions about cell-type composition, reference signatures, or linear mixing that frequently fail to capture the complexity and plasticity of malignant states. As a result, bulk RNA-seq continues to lack the necessary resolution to define cancer cell lineage programs, intertumoral heterogeneity, and applicability of experimental models.

Single-cell RNA sequencing (scRNA-seq) offers the resolution necessary to isolate malignant transcriptional states and disentangle cancer cell-specific programs from surrounding microenvironmental signals. This promise has motivated the construction of pan-cancer scRNA-seq atlases.^5–10^ However, these efforts face substantial challenges that limit their biological utility and adoption.^11,12^ Publicly available scRNA-seq datasets vary widely in sequencing depth, technical quality, preprocessing strategies, and annotation accuracy, particularly across diverse cancer types where automated cell-type labeling performs inconsistently. Further, and perhaps most importantly, atlas efforts tend to prioritize maximal data aggregation, producing increasingly large and computationally unwieldy resources that trade biological coherence for scale. These atlases require substantial computational infrastructure, lack clear criteria for malignant cell representation, and offer limited interpretability for downstream translational applications. Consequently, despite containing an enormous amount of data, these resources are infrequently used as practical references for tumor comparison, model evaluation, or hypothesis generation.

Addressing these challenges is essential for the broader use of pan-cancer data, particularly as large-scale functional genomics efforts continue to expand. There is now an enormous body of high-throughput gene dependency data using cancer cell lines.^13–18^ These datasets provide an unparalleled opportunity to link transcriptional state to functional vulnerabilities. However, dependency models that apply these data to cancers *in vivo* are currently trained on bulk RNA-seq profiles from cell lines and human tumors, inheriting the same limitations that complicate bulk RNA-seq-based tumor comparisons. Integration between cell line bulk RNA-seq and tumor scRNA-seq remains challenging and often unreliable beyond coarse gene set correlations, underscoring the need for scRNA-seq–to–scRNA-seq comparisons to ultimately link high-throughput cancer cell line data to human tumors.

Together, these limitations motivated the development of a high-quality, biologically coherent, single-cell reference encompassing both human tumor and cancer cell line transcriptional programs. We applied strict filtering and standardized processing to publicly available scRNA-seq tumor datasets to assemble a pan-cancer atlas (scTumor Atlas) containing more than 135,424 high-quality malignant cells from 499 samples spanning 36 adult and pediatric cancer types. Stringent quality control and Mahalanobis distance-based cell selection produced a balanced and biologically coherent dataset suitable for cross-cancer comparisons. Pathway scoring identified tumor-specific transcriptional programs and highlighted cancers with highly consistent biological identities across datasets. The validity of these programs was supported by pathway-based agreement with expression patterns observed in established TCGA bulk RNA-seq cohorts. For each tumor type, cancer cell lines were evaluated against the scTumor Atlas-derived signatures, generating models that revealed preserved malignant programs and highlighted transcriptional divergence between many cell lines and cancer of origin. Finally, we leveraged DepMap CRISPR screening data to construct a regression model used to predict gene dependency from scRNA-seq data, and we demonstrated ease of use and applicability in a personalized manner with patient-derived scRNA-seq data in our laboratory. Overall, this atlas provides a compact and biologically grounded reference for comparing tumors, interpreting pathway structure, and assessing applicability of cancer cell line-based data.

## Results

### Construction of a High-Quality Pan-Cancer Single-Cell Tumor Atlas (scTumor Atlas)

To establish a foundation for cross-cancer analysis, we aggregated publicly available scRNA-seq datasets to construct the scTumor Atlas, comprising 36 adult and pediatric malignancies across all major anatomical systems (**Fig**. **1A****, Supplementary Table 1**). To address known variability in sequencing platforms and read depth, we applied a uniform, stringent quality control pipeline (**Fig**. **1B**). Cells with <5,000 counts or >10% mitochondrial transcripts were excluded, as were samples containing <200 cells. Doublets were identified and removed with Scrublet.^19^ Malignant populations were then identified based on prior annotation or through manual annotation using lineage markers, inferred copy-number variation, and transcriptional proximity to expected tumor states (**Extended Data Fig. 1**).

**Fig. 1.**
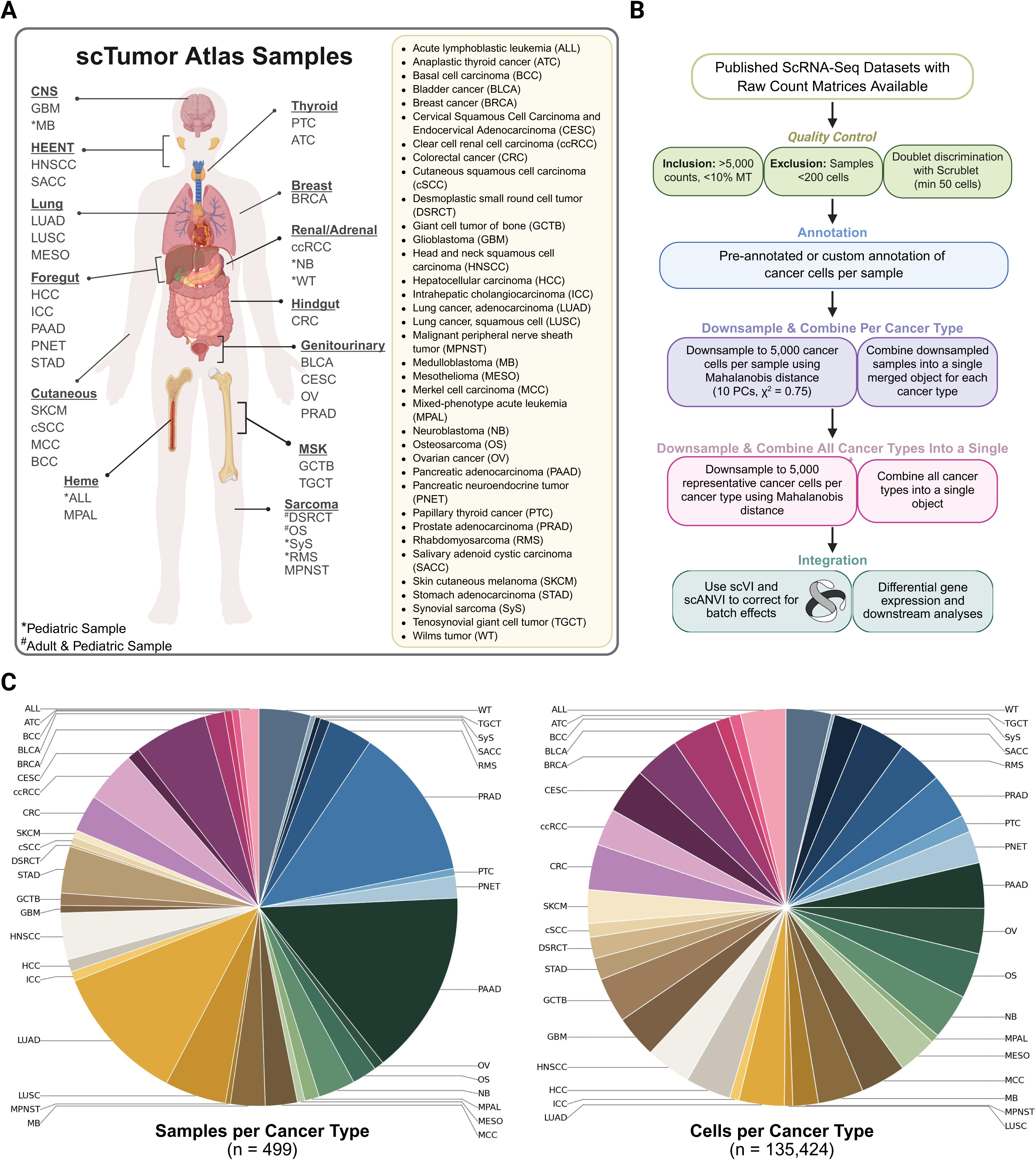
Construction of a high-quality, pan-cancer, single-cell tumor atlas (scTumor Atlas). (**A**) Representative anatomical schematic illustrating the distribution of the 36 adult and pediatric cancers included in the scTumor Atlas across major organ systems. (**B**) ScTumor Atlas construction workflow. Raw count matrices from published scRNA-seq datasets were uniformly reprocessed. Quality control excluded cells with <5,000 UMIs, >10% mitochondrial content, samples with <200 cells, and doublets identified using Scrublet. Malignant cells were annotated for each sample. Individual samples were downsampled to 5,000 representative malignant cells using Mahalanobis distances from the centroid in principal component space. These samples were then combined by cancer type following Harmony integration. A second atlas-level downsampling step retained up to 5,000 representative cancer cells per cancer type for downstream cross-cancer analyses. (**C**) ScTumor Atlas composition. Pie chart distribution of the 499 tumor samples (left) and 135,424 malignant cells (right) comprising the scTumor Atlas.

To prevent individual datasets from disproportionately influencing downstream analyses, we developed a two-step downsampling framework. For each cancer type, up to 5,000 representative malignant cells were selected using Mahalanobis distance from the centroid in principal component space, capturing consistent transcriptional features while equilibrating dataset representation. All subsequent datasets within a cancer type were merged and integrated using Harmony,^20^ followed by a second atlas-level downsampling to 5,000 representative cells per cancer type. After filtering and annotation, the atlas comprised 499 samples and 135,424 high-quality cancer cells (**Fig**. **1C**).

### Integrated Analysis Demonstrates Lineage as a Major Axis of Cross-Cancer Organization

To validate the global organization of the scTumor Atlas, we used scVI^21^ and scANVI^22^ to generate an integrated latent space that minimized residual technical structure and preserved consistent biological patterns across cancer types. Incorporating partial annotations with scANVI further sharpened boundaries between malignant states and improved resolution of cancer-level differences (**Fig**. **2A**). As a brief overview, epithelial tumors demonstrated canonical enrichment of *EPCAM* and *KRT8*, mesenchymal tumors exhibited elevated *COL1A1* expression, hematologic cancers expressed immune lineage markers including *CD69*, and neuroendocrine tumors demonstrated strong *TSPAN7* and *CHGA* expression **(Fig. 2B, Supplementary Table 2).** To assess potential technical contributions to the observed structure, sequencing depth and library complexity were evaluated across cancer types. Total counts and detected genes fell within a comparable range across tumor types (**Fig. 2C**). Heatmap analysis of differential expressed genes highlighted cancer type-specific transcriptional programs. (**Fig**. **2D**). Dot plot visualization of canonical markers further confirmed preservation of cancer type-specific expression patterns across the atlas (**Fig. 2E, Supplementary Table 2**). For example, mesothelioma (MESO) cells showed high expression of *MSLN*, whereas melanoma (SKCM) cells showed high expression of *MLANA*. Together, these results suggest that data integration retained biologically meaningful distinctions without blurring tumor-specific transcriptional identities.

**Fig. 2.**
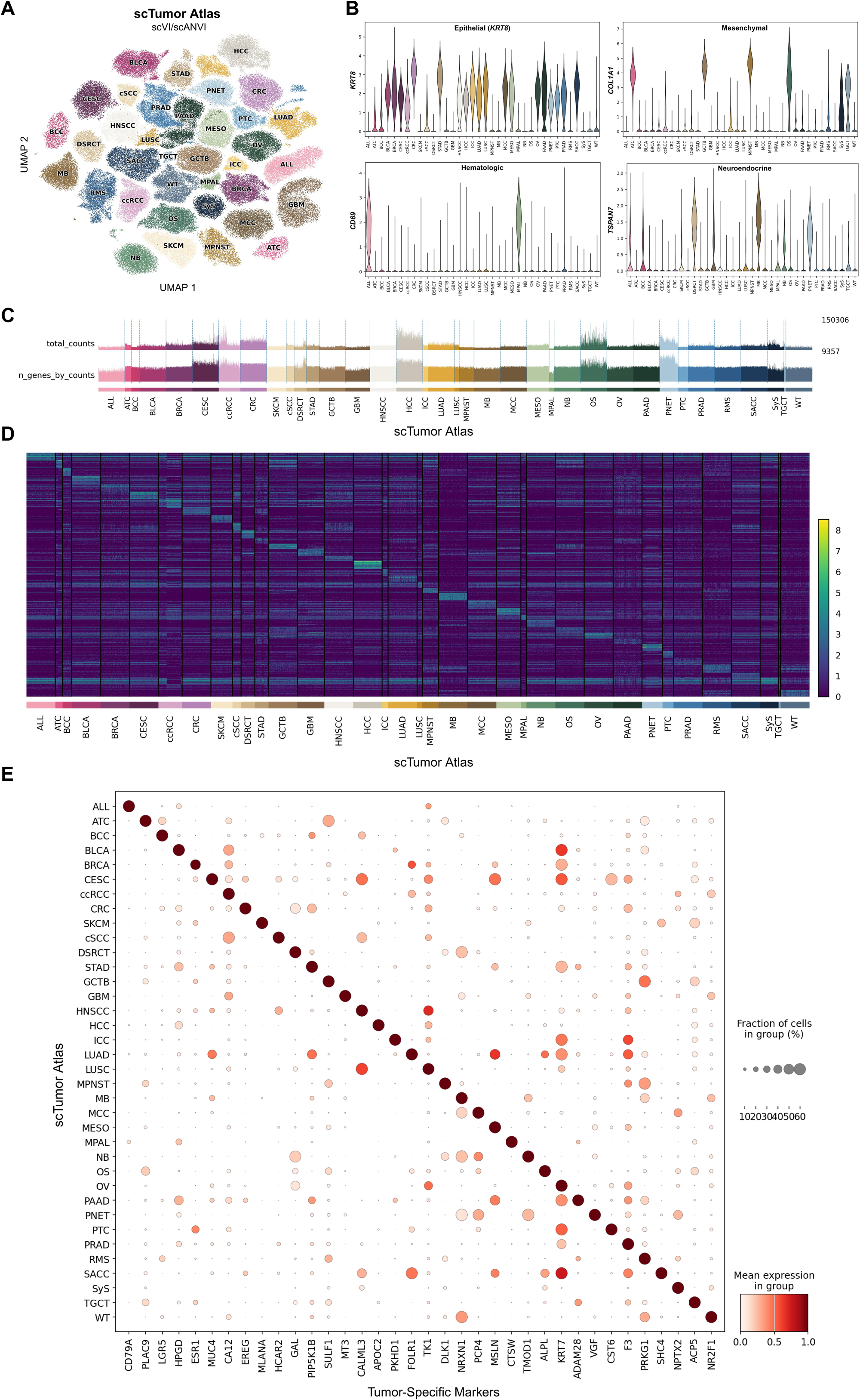
Preservation of cancer-type identity in the integrated single-cell Tumor (scTumor) Atlas. (**A**) Uniform manifold approximation and projection (UMAP) of cancer types in the scTumor Atlas after integration with scVI and refinement with scANVI. (**B**) Expression of canonical lineage-associated markers across the atlas, including *KRT8* (epithelial), *COL1A1* (mesenchymal), *CD69* (hematologic), and *TSPAN7* (neuroendocrine). (**C**) Track plot highlighting the distribution of raw library metrics, including total UMI counts and number of detected genes, across cancer types in the scTumor Atlas. (**D**) Heatmap of the top 20 differentially expressed genes by tumor type. (**E**) Dot plot of canonical marker gene expression across cancer types.

### Cross-Cancer Pathway Analysis Identifies Conserved and Tumor-Specific Biological Programs

To validate preservation of lineage-specific tumor biology within the atlas, gene set enrichment analysis (GSEA) was performed. A cross-cancer heatmap of normalized enrichment scores revealed a highly coherent organization of pathway activity, with cancer types organizing into groups defined by metabolic, proliferative, immune, and developmental programs (**Fig**. **3A****, Extended Data Fig. 2, Supplementary Table 3**). Pathway-level analyses further reinforced lineage-expected biological activity across tumor types. Oxidative phosphorylation was strongly enriched in lung squamous carcinoma (LUSC) but minimally present in acute lymphoblastic leukemia (ALL), consistent with prior reports of elevated oxidative phosphorylation activity in lung cancer (**Fig**. **3B**).^23,24^ MYC target gene sets were enriched in ALL and depleted in pancreatic adenocarcinoma (PAAD) (**Fig. 3C**).^25^ Hormone response pathways reflected expected tissue specificity, with androgen signaling enrichment in prostate cancer (PRAD) and estrogen signaling enrichment in breast cancer (BRCA) (**Fig**. **3D**). Upregulated KRAS signaling in PAAD (**Fig**. **3E**) and elevated epithelial to mesenchymal transition (EMT) signatures in sarcomas and anaplastic thyroid cancer (ATC) further support the biological fidelity of the tumor atlas (**Fig. 3F**).^26–30^

**Fig. 3.**
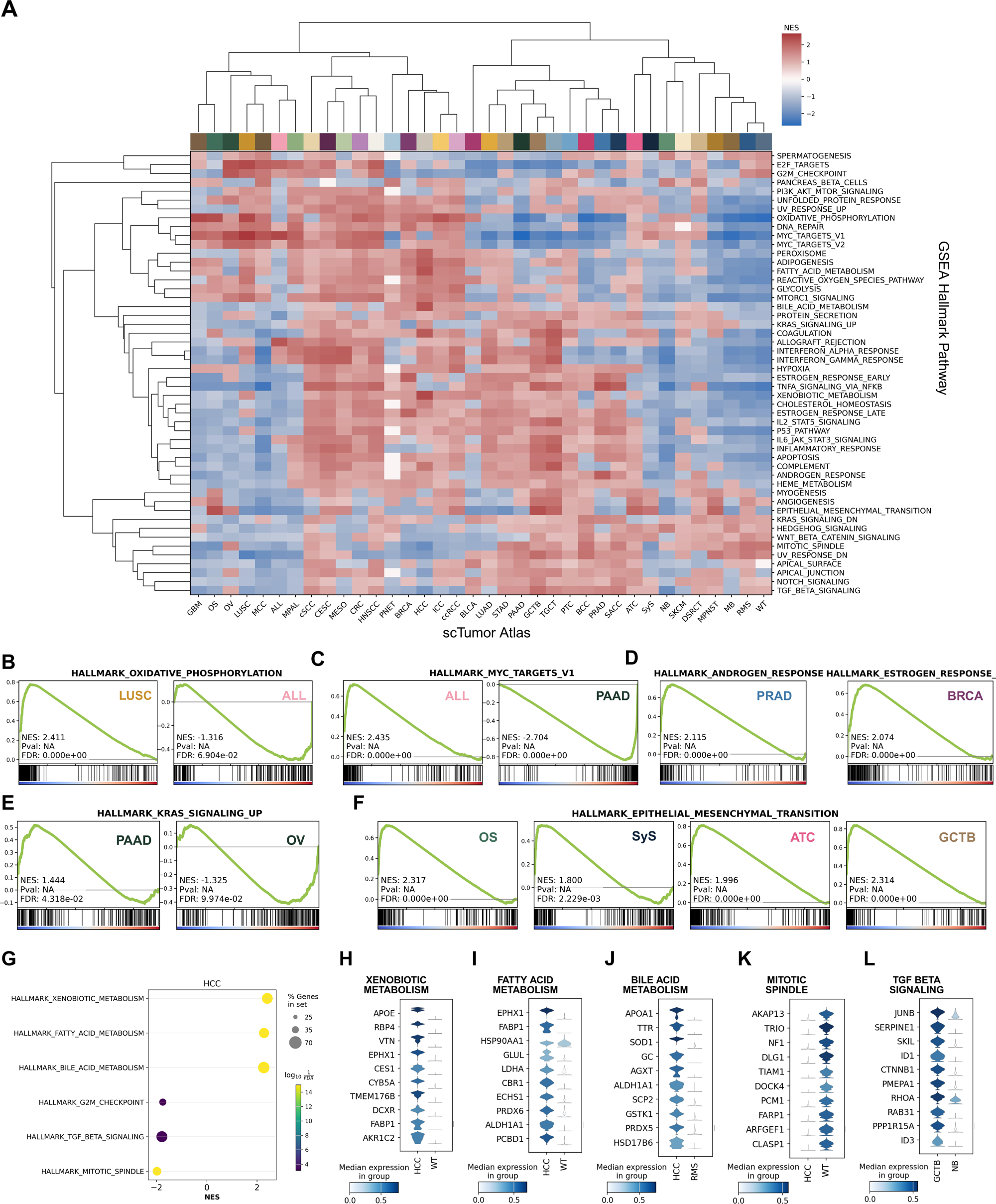
Cross-Cancer Pathway Activity and Lineage-Specific Biological Programs. **(A)** Heatmap of Gene Set Enrichment Analysis (GSEA) normalized enrichment scores (NES) for Hallmark gene sets by scTumor Atlas cancer type. **(B)** GSEA enrichment plots of oxidative phosphorylation enrichment in lung squamous cell carcinoma (LUSC) and acute lymphoblastic leukemia (ALL). **(C)** Enrichment plots illustrating MYC target activity in ALL and pancreatic adenocarcinoma (PAAD). **(D)** GSEA plots demonstrating the enrichment of androgen response pathways in prostate cancer (PRAD) and estrogen response pathways in breast cancer (BRCA). **(E)** KRAS signaling GSEA enrichment plots in PAAD and ovarian carcinoma (OV). (**F**) GSEA enrichment plots demonstrating the enrichment pattern of epithelial–mesenchymal transition (EMT) pathway across mesenchymal tumors including osteosarcoma (OS), synovial sarcoma (SyS), anaplastic thyroid carcinoma (ATC), and giant cell tumor of the bone (GCTB). (**G**) Dot plot illustrating the top 3 upregulated and downregulated GSEA Hallmark pathways in hepatocellular carcinoma (HCC). **(H)** Violin plots for xenobiotic metabolism genes in HCC compared with Wilms tumor (WT). (**I**) Violin plots for fatty acid metabolism genes in HCC compared with WT. **(J)** Violin plots highlighting the expression of various genes in bile acid metabolism (HCC versus rhabdomyosarcoma (RMS)). **(K)** Violin plots for mitotic spindle genes in HCC compared with WT. **(L)** Violin plots for TGF-β signaling genes in GCTB compared with neuroblastoma (NB).

Hepatocellular carcinoma (HCC) demonstrated expected enrichment of xenobiotic metabolism, fatty acid metabolism, bile acid metabolism, and related metabolic programs (**Fig**. **3G**). Violin plots confirmed uniform expression of key metabolic genes in these pathways, including *APOE*, *CES1, FABP1, ALDH1A1*, and *SCP2* (**Fig**. **3H-K**), demonstrating coordinated elevation across pathway rather than enrichment driven by any single transcript. The TGF-β pathway demonstrated cancer-specific activation in GCTB, reflecting the established abundance of TGF-β isoforms released during tumor-induced osteoclastic bone resorption and representing an expected biological benchmark (**Fig**. **3L**).^31–33^ Together, these results suggest both shared and lineage-specific biological programs, confirming the biological fidelity of the atlas and potential suitability for downstream modeling.

### Bulk RNA Sequencing Recapitulates Lineage Structure and Confirms Concordance with Single-Cell Signatures

To benchmark malignant programs identified in the scTumor Atlas against independent bulk tumor data, we analyzed TCGA bulk RNA-sequencing (bulk RNA-seq) data from 8,246 matching tumor types. Bulk samples formed well-defined clusters by cancer type, with clear separation of epithelial, mesenchymal, and hematologic lineages (**Fig**. **4A**). Previously identified lineage markers defined in the scTumor Atlas (**Fig. 2E**) were also expressed in the corresponding TCGA cancer types (**Fig**. **4B**), demonstrating reasonable concordance between single-cell–derived signatures and established bulk tumor profiles.

**Fig. 4.**
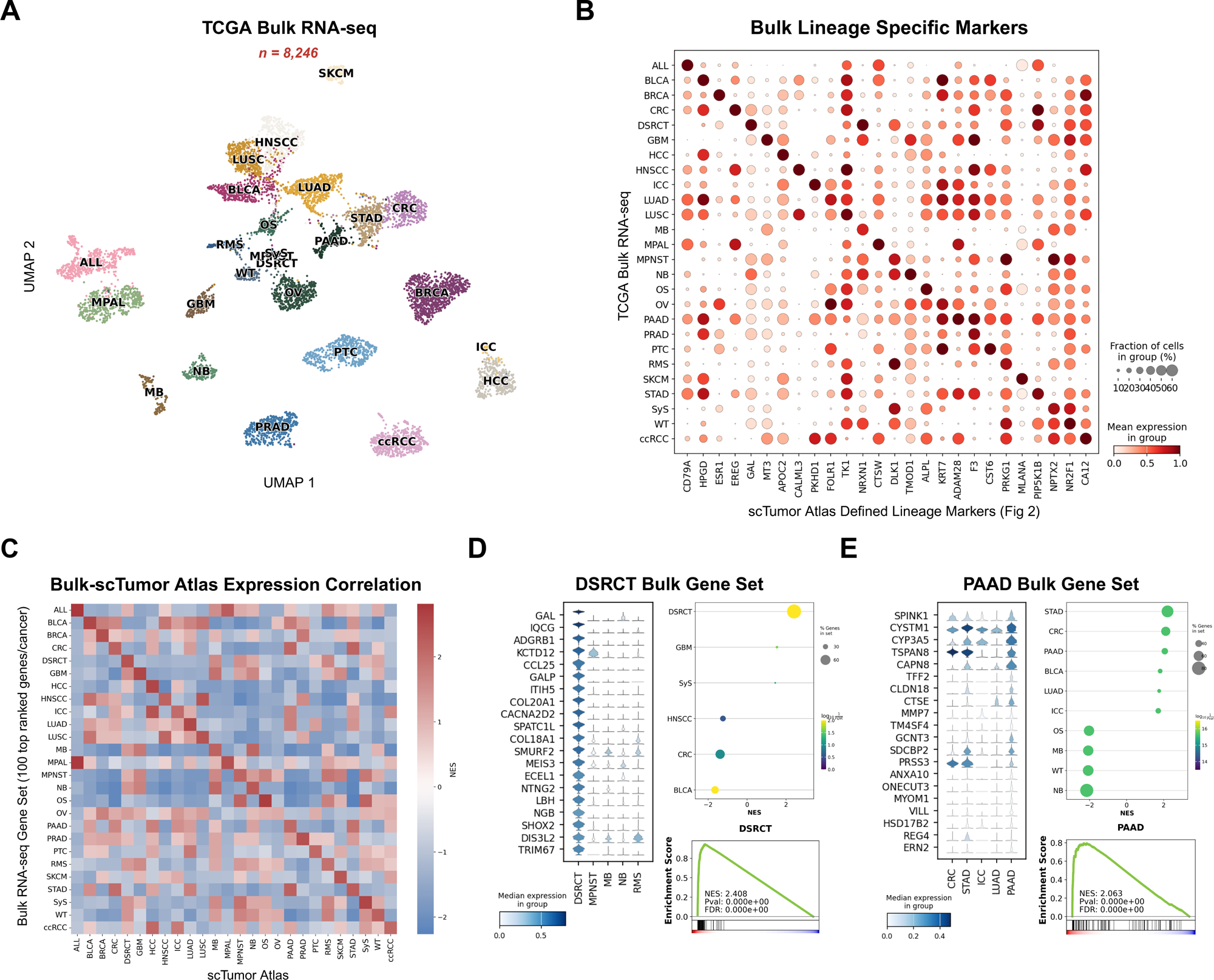
Bulk RNA-seq Profiles and Correspondence with Single-Cell Cancer Types. **(A)** UMAP illustrating 8,246 The Cancer Genome Atlas (TCGA) bulk RNA-seq samples colored by cancer type. **(B)** Dot plot demonstrating expression of the same genes highlighted in Fig. 2E across TCGA samples. **(C)** Heatmap of GSEA normalized enrichment scores (NES) derived from the top 100 differentially expressed genes in bulk RNA-seq (bulk gene sets), evaluated against differentially expressed genes across scTumor Atlas cancer types. **(D)** Violin plot of desmoplastic small round cel tumor (DSRCT) bulk gene set expression in DSRCT, malignant peripheral nerve sheath tumors (MPNST), medulloblastoma (MB), neuroblastoma (NB), and rhabdomyosarcoma (RMS) (left), and GSEA plots testing the DSRCT bulk-derived gene set against scTumor Atlas cancer types (right). **(E)** Violin plot displaying expression of the PAAD bulk gene set in colorectal cancer (CRC), gastric adenocarcinoma (STAD), intrahepatic cholangiocarcinoma (ICC), lung adenocarcinoma (LUAD), and PAAD (left), and GSEA enrichment plots testing the PAAD bulk-derived signature against scTumor Atlas cancer types (right).

To further assess preservation of native transcriptomic profiles, we derived bulk RNA-seq gene sets from the top 100 ranked genes per cancer type and performed GSEA on scTumor Atlas profiles (**Fig**. **4C****, Supplementary Table 4**). This analysis revealed a high degree of correspondence between bulk RNA-seq and single-cell profiles, with the notable exception of ovarian cancer (OV). Several tumor types also exhibited secondary associations with biologically related cancers, such as mixed phenotype acute leukemia (MPAL) and acute lymphoblastic leukemia (ALL). Next, we were interested in whether more homogeneous tumors in the scTumor atlas would show higher enrichment of bulk gene sets than more heterogeneous tumors. Focusing on representative examples, desmoplastic small round cell tumor (DSRCT) showed strong concordance between single-cell and bulk profiles, consistent with the high proportion of relatively homogenous malignant cells in these specimens (**Fig**. **4D**). In contrast, pancreatic adenocarcinoma (PAAD) displayed greater divergence between modalities, reflecting its pronounced cellular heterogeneity and complex transcriptional landscape (**Fig**. **4E**).

Together, these analyses demonstrate that malignant programs identified in the scTumor Atlas align reasonably well with established bulk RNA-seq signatures, providing independent validation of cancer type-specific gene expression patterns.

### Assessment of Cancer Cell Line Fidelity Using Bulk, Single-Cell, and scTumor Atlas Projections

Given the widespread use of cancer cell lines (CCLs) as in vitro tumor models, we evaluated how scTumor Atlas–defined malignant programs are reflected in CCLs derived from corresponding tumor types, leveraging publicly available bulk and single-cell RNA sequencing datasets for benchmarking and modeling. While bulk RNA-seq of CCLs is widely used, scRNA-seq studies of CCLs are smaller and not as well incorporated into CCL workflows; therefore, we first assessed whether existing scRNA-seq data from CCLs reflected established bulk RNA-seq patterns for the same CCLs. Across the 206 CCLs evaluated, scRNA-seq (scCCL) UMAPs formed well-defined clusters at the individual cell line level compared to bulk RNA-seq (bCCL). In contrast, cancer type of origin was less clearly resolved in both scCCL and bCCL expression profiles, indicating that major lineage-associated transcriptional features were not readily captured by either sequencing modality (**Fig. 5A-B**). To directly compare transcriptional structure across modalities, we performed pairwise comparisons using canonical correlation analysis (CCA), which demonstrated excellent correspondence between bulk CCL profiles and their pseudobulk scCCL counterparts (**Fig**. **5C-D**). At the individual tumor type level, pancreatic adenocarcinoma (PAAD) CCLs exhibited stable transcriptional signatures across bulk and single-cell datasets, supporting preservation of core malignant programs despite modality-specific differences (**Fig**. **5E****, Extended Data Fig. 3**). These data confirm that the transcriptomic profile of CCLs captured by scRNA-seq is comparable to established bulk RNA-seq data from the same CCLs.

**Fig. 5.**
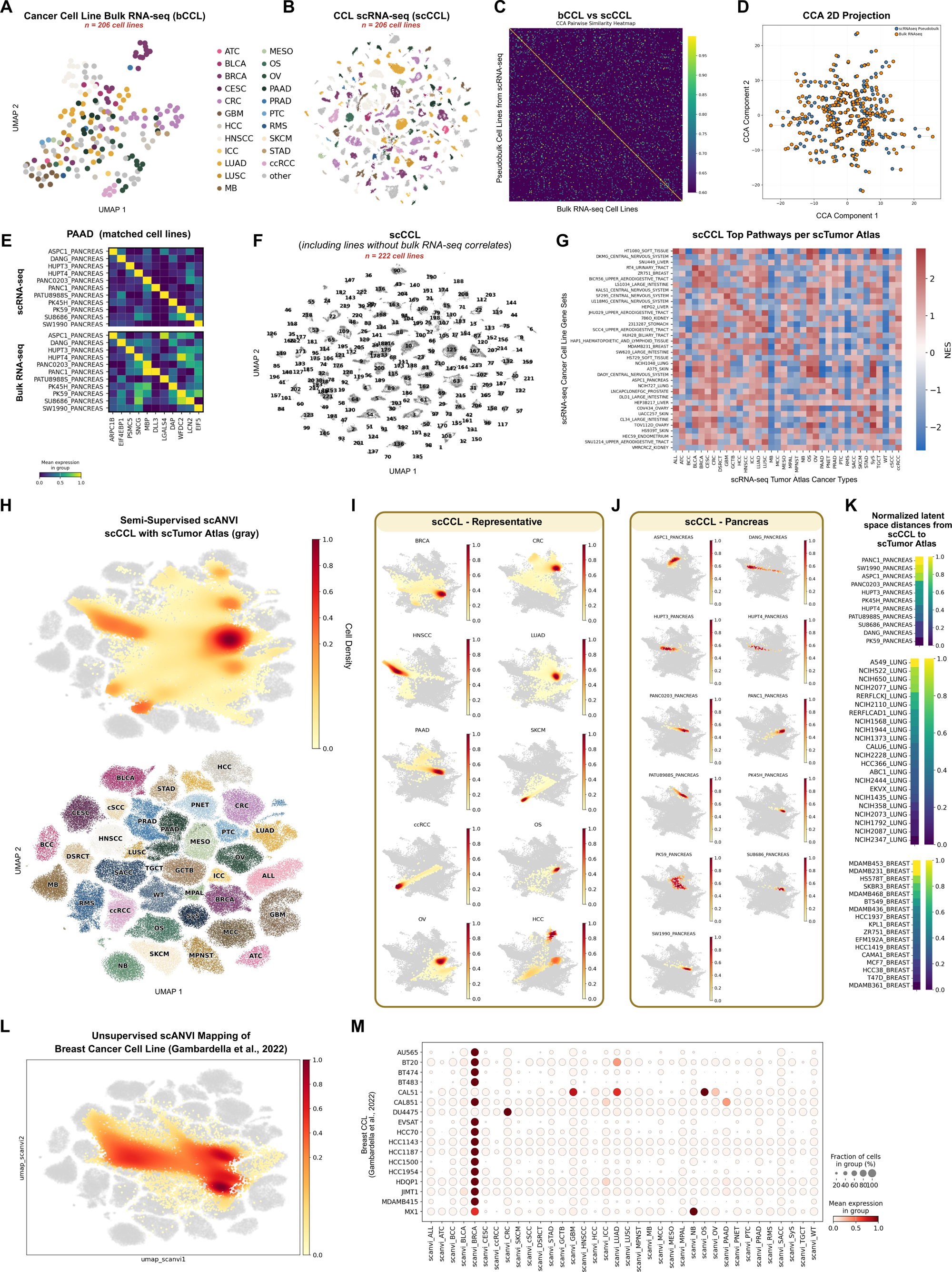
Evaluating Cancer Cell Line (CCL) Fidelity Using Bulk, Single-Cell, and Atlas Projections. (**A**) Uniform manifold approximation and projection (UMAP) of bulk RNA sequencing (RNA-seq) profiles from 206 publicly available cancer cell lines (bCCL). Each dot represents an individual bulk RNA-seq sample. (**B**) Single cell RNA-seq (scRNA-seq) UMAP of 206 corresponding cancer cell lines (scCCL). (**C**) Heatmap demonstrating canonical correlation analysis (CCA) or pairwise similarities between bCCL data and pseudobulked scCCL data. (**D**) A two-dimensional rendering of CCA pairwise similarity analysis in (C), colored by sequencing modality. Dashed lines connect paired cancer cell lines across modalities (scRNA-seq and bulk RNA-seq). (**E**) Heatmap illustrating gene expression across matched scRNA-seq and bulk RNA-seq of pancreatic adenocarcinoma (PAAD) cell lines. (**F**) UMAP of 222 publicly available single-cell RNA sequencing (scRNA-seq) cancer cell lines, highlighting distinct populations. This single-cell cancer cell line (scCCL) dataset includes 16 cell lines excluded in (B) due to the absence of a bulk RNA-seq correlate. (**G**) Heatmap of normalized enrichment scores (NES) of curated scCCL gene sets (top 100 differentially expressed genes per scCCL sample) in the scTumor Atlas. **(H)** Projection of scCCL profiles onto the scTumor Atlas scANVI UMAP using normalized Euclidean distances in shared latent space. **(I)** Representative scCCL scANVI UMAP projections. **(J)** Projections of individual pancreatic adenocarcinoma (PAAD) cell lines. **(K)** Heatmaps illustrating normalized latent-space distances of scCCLs (pancreas, lung, and breast cancer) to their corresponding primary tumor centroids in the scTumor Atlas. Lower distance (blue) represents higher cell line fidelity to the scTumor Atlas centroid. (**L**) Application of the pretrained scTumor Atlas scANVI model to previously unseen breast cancer cell line scRNA-seq datasets,^34^ enabling nearest-neighbor projection onto the reference UMAP. (**M**) Dot plot of scTumor Atlas cancer-type prediction probabilities for previously unseen breast cancer cell lines using the trained scANVI model.

To assess concordance between scCCL and the scTumor Atlas, we defined gene sets using the top 100 differentially expressed genes from each CCL and performed GSEA across scTumor Atlas cancer types. In this framework, CCL-derived gene signatures were treated as gene sets, and scTumor Atlas cancer types were used as the enrichment background, analogous to Hallmark pathway analysis (**Fig**. **5F-G****, Supplementary Table 5**). Normalized enrichment scores from this analysis demonstrated high concordance between certain CCLs and cancer of origin, but this observation was not consistent (**Fig**. **5F-G**). To more directly assess CCL transcriptional profiles within the scTumor Atlas framework, we trained the scTumor Atlas scANVI model on the scCCL data. This resulted in a co-embedding in which cancer-type programs conserved in CCL propagation could be identified. As a visual aid, nearest neighbor analysis between CCLs and primary cells in the shared latent space was used to project approximate locations of CCL cells onto the original scTumor Atlas UMAP. The projection aligned many cancer-type-specific programs between primary tumors and CCLs (**Fig**. **5H-I**); however, considerable diversity remained among CCLs within a cancer type (**Fig. 5J**). For example, centroid-based Euclidean distance analysis revealed that selected PAAD cell lines, including PK59 and DANG, exhibited short normalized latent-space distances, corresponding to greater transcriptional similarity to primary PAAD tumor centroids, whereas PANC1 and SW1990 showed substantially greater distances, indicating divergence from primary tumor transcriptional states (**Fig. 5J-K, Extended Data Fig. 4**). These approximate measures suggest potential utility in evaluating the translational potential of high-throughput CCL-based experiments beyond categorization of cancer type of origin.

To evaluate whether the model could be used to predict cancer type of origin for a given CCL, we applied our scANVI model in an unsupervised manner to a publicly available breast CCL scRNA-seq dataset.^34^ We observed accurate prediction of cancer type of origin in 14 of the 17 breast CCLs (**Fig**. **5L-M**). Together, these results demonstrate that scTumor Atlas-derived transcriptional signatures may provide a robust and interpretable framework for evaluating CCL similarity to its cancer type of origin *in vivo*. This approach supports more informed selection of experimental models and enhances the clinical relevance of preclinical predictions.

### Single-cell expression profiles can be used to predict CRISPR gene dependencies across cancer types

To model gene-level dependencies from transcriptional state, we trained regularized regression models that predict Chronos gene-effect scores directly from matched scCCL expression profiles, adapting a method originally developed for bulk RNA-seq data.^17,35–37^ In brief, we fit ElasticNet regression models independently for each target gene, using cross-validation to balance sparsity and model stability. Only genes exhibiting sufficient dependency signal across cell lines (Chronos score < −1) were included to ensure robust model estimation. Within the CCL cohort, expression features were standardized and used to model the relationship between transcriptional programs and gene essentiality. Model performance was evaluated by cross-validation, and high-confidence models were defined by a stringent performance threshold (coefficient of determination ≥ 0.5), yielding a subset of genes for which expression reliably predicted functional dependency. The learned regression coefficients were then applied to standardized single-cell expression profiles from the scTumor Atlas to generate predicted gene-effect scores (PGES) at single-cell resolution. PGES values were subsequently normalized across cells to enable cross-dependency comparisons. This framework enabled systematic inference of cancer-specific genetic vulnerabilities directly from single-cell transcriptional states while maintaining alignment with large-scale functional genomics screens (**Fig**. **6A****, Supplementary Table 6**).^35,36^

**Fig. 6.**
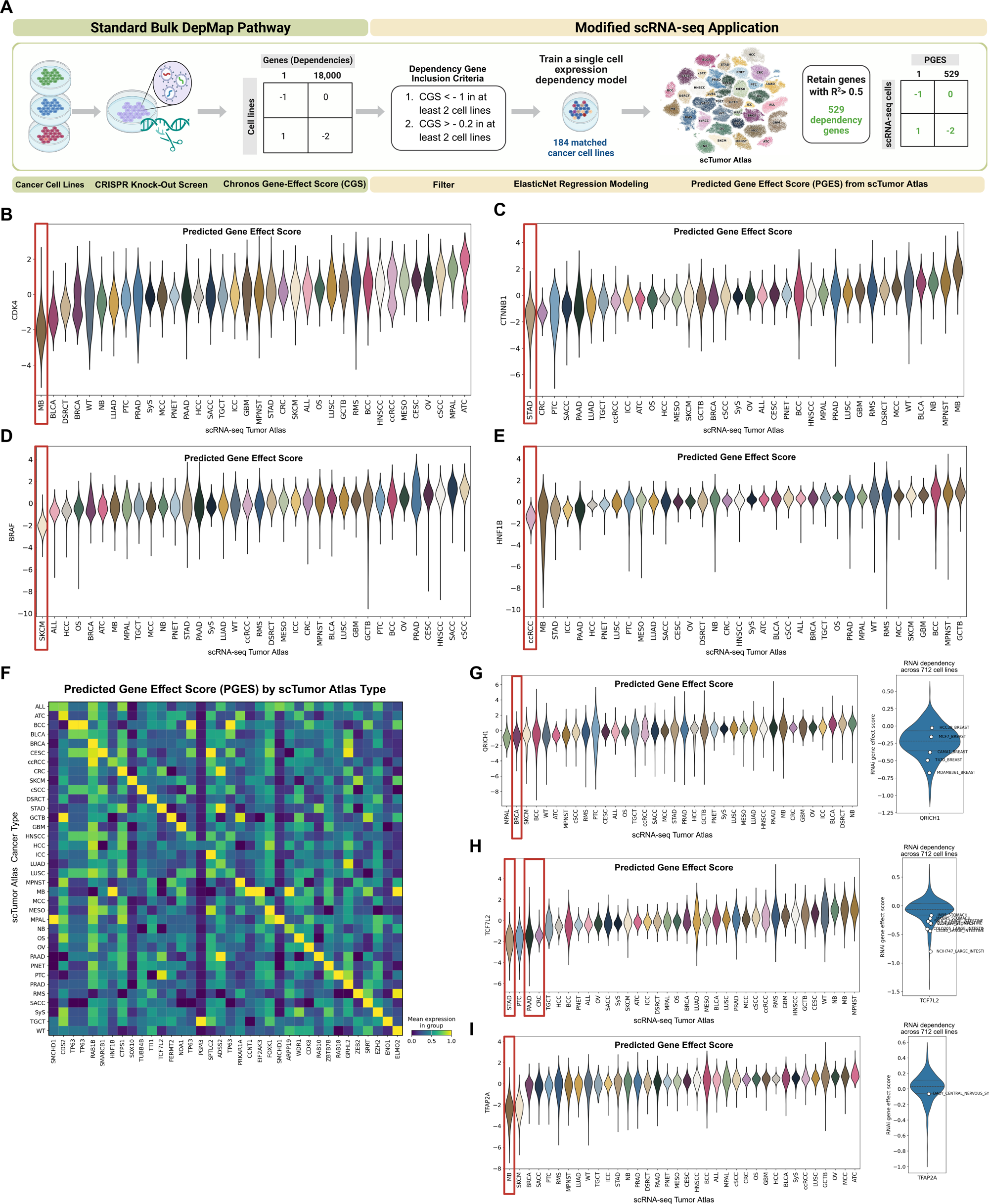
Single-Cell Gene Expression Enables Prediction of DepMap CRISPR Screen–Derived Gene Dependencies. (**A**) Overview of the scRNA-seq dependency-modeling workflow. (**B–E**) Predicted gene effect scores (PGES) for *CDK4, CTNNB1, BRAF*, and *HNF1B* across scTumor Atlas cancer types. More negative values indicate greater cell dependency on the tested gene. (**F**) Heatmap of the top PGES by scTumor Atlas cancer type. (**G-I**) PGES of previously unexplored dependency genes across scTumor Atlas cancer types (left) and corresponding orthogonal RNA interference (RNAi) dependency scores in matched cancer cell lines (right). Cell lines with the highest fidelity to the scTumor centroid (lowest normalized latent-space distance) are shown as individual points in the RNAi plots.

These regression models recapitulated established lineage-specific dependencies, such as *CDK4* dependence in medulloblastoma (MB), *CTNNB1* in gastric cancer (STAD), *BRAF* in melanoma (SKCM), and *HNF1B* dependence in clear-cell renal carcinoma (ccRCC) (**Fig**. **6B-E**).^38–41^ Across the atlas, predicted dependencies organized into distinct, cancer-specific patterns, with each tumor type exhibiting a distinct set of vulnerabilities (**Fig**. **6F**). Beyond recapitulating known targets, this framework identified putative novel vulnerabilities, including *QRICH1* in breast cancer (BRCA), *TCF7L2* in gastric (STAD), pancreatic (PAAD), and colorectal cancer (CRC), and *TFAP2A* in medulloblastoma (MB) (**Fig**. **6G-I** **(left panel), Extended Data Fig. 5**).^42–45^

Independent validation of CRISPR-derived dependencies was performed using DepMap RNA interference (RNAi) data with representative cell lines,^17,37,42^ previously selected based on normalized Euclidean distance to *in vivo* cancer type centroids in the shared latent space as a measure of concordance with primary tumor states (**Fig**. **6G-I** **(right panel), Fig. 5K**). Together, these results demonstrate a degree of orthogonal validation for predicted dependencies. By enabling inference of functional vulnerabilities directly from scRNA-seq data, this approach provides a scalable framework for identifying therapeutic targets across diverse cancer types, with the potential to evolve into a personalized approach.

### Personalized model application to scRNA-seq data in a rare tumor

To demonstrate ease of use and potential translational application of atlas-based dependency prediction, we projected an internally sequenced primary retroperitoneal leiomyosarcoma (RPLMS) into the scTumor Atlas latent space. Manual annotation of scRNA-seq data demonstrated malignant cells expressing canonical smooth muscle markers (**Fig**. **7A-B**).

**Fig. 7.**
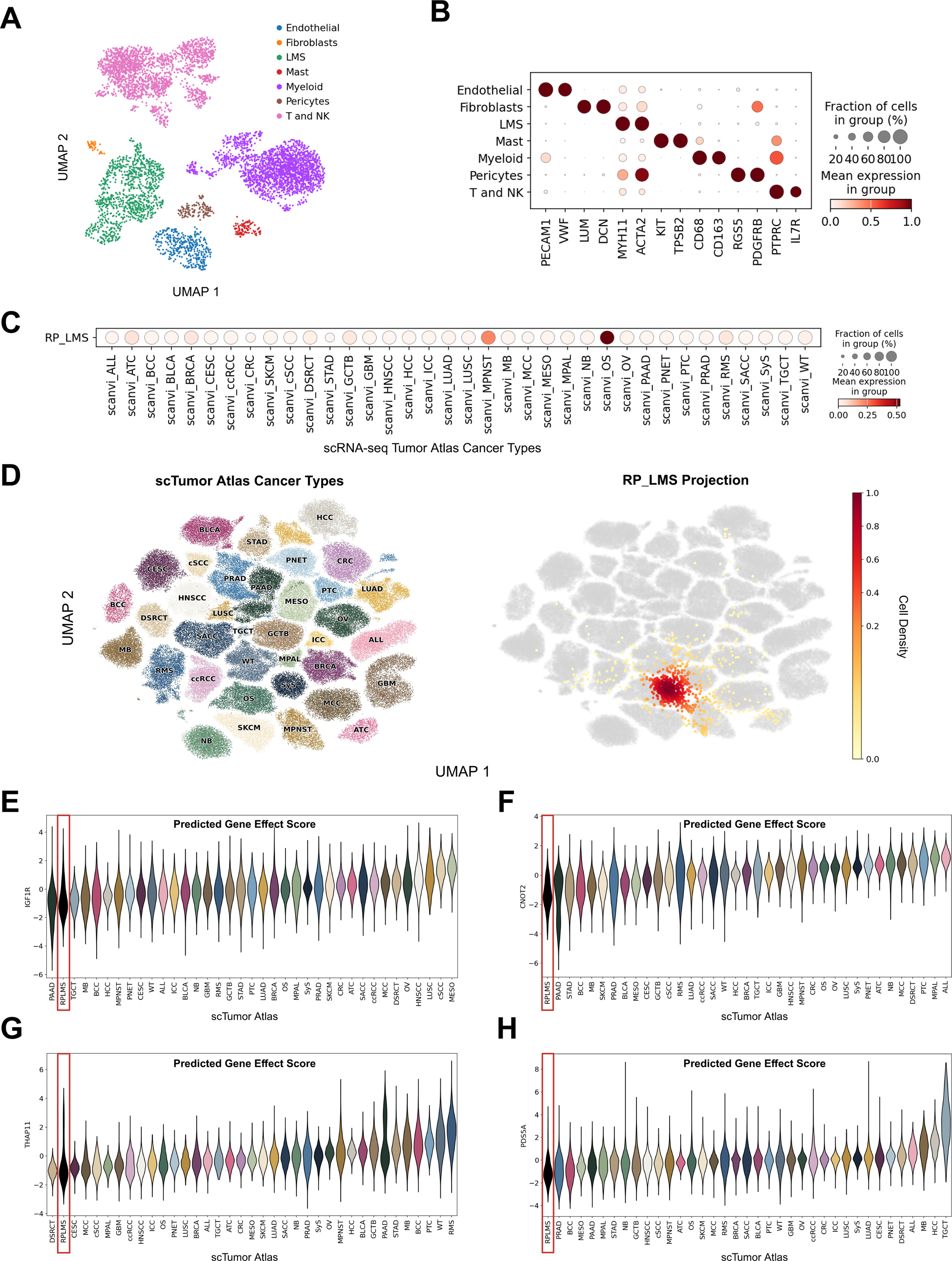
Validation of Single-Cell-Based CRISPR Dependency Predictions in Primary Retroperitoneal Leiomyosarcoma (RPLMS). **(A)** UMAP visualization of major cell types in RPLMS, including immune, stromal, endothelial, and tumor compartments in an internally generated RPLMS dataset. **(B)** Dot plot of canonical markers of each manually annotated cell type. (**C**) Dot plot showing scTumor Atlas cancer type prediction probabilities for previously unseen RPLMS cells derived from the retrained scTumor scANVI model. (**D**) Nearest-neighbor projection of RPLMS cels onto the scTumor scANVI UMAP**. (E–H)** Predicted gene effect scores (PGES) for *IGF1R*, *CNOT2*, *THAP1*, and *PDS5A* across scTumor Atlas cancer types following RPLMS integration.

Per-cell cancer type assignment probabilities were inferred using scANVI and used to assess relative similarity to reference cancer types from the scTumor Atlas, summarized using dot plots of mean label probabilities. This comparative similarity scoring demonstrated rough alignment of this RPLMS with other sarcomas (**Fig**. **7C-D**). Application of the developed regression models identified candidate gene dependencies in this RPLMS, including *IGF1R*, *CNOT2*, *THAP11,* and *PDS5A*, implicating pathways involved in cell survival, transcriptional regulation, mRNA stability, and mitosis. (**Fig**. **7E–H**). IGF1R is known to be upregulated in leiomyosarcoma and has been targeted in clinical trials of leiomyosarcoma patients.^43–46^ These candidate vulnerabilities represent actionable hypotheses for future functional validation. Together, these findings demonstrate that the scTumor Atlas framework can be used for functional dependency inference from scRNA-seq data of a single patient tumor. These findings highlight the real-world translational potential of this workflow, particularly for rare cancers.

## Discussion

In this study, we present the scTumor Atlas, a compact and biologically coherent pan-cancer single-cell reference spanning 36 adult and pediatric malignancies that links tumor identity to cancer cell line evaluation and functional dependency inference. By prioritizing stringent quality control, balanced inclusion, and representative malignant state selection over maximal data aggregation, the atlas preserves lineage-specific transcriptional programs while remaining computationally lightweight and broadly deployable. The scTumor Atlas enables direct scRNA-seq to scRNA-seq comparison between human tumors and cancer cell lines, supporting inference of CRISPR-derived gene dependencies at single-cell resolution. This framework addresses a longstanding barrier to translating large-scale functional genomics data into biologically and clinically meaningful contexts.

This work builds on foundational pan-cancer single-cell efforts, including the Curated Cancer Cell Atlas^5,8^ and CancerSCEM,^7,47^ which established the feasibility and value of large-scale tumor profiling, as well as studies defining recurrent malignant programs across cancers.^48,49^ In contrast to atlas efforts emphasizing exhaustive aggregation, our approach centers on representative malignant transcriptional states, yielding a reference that is both interpretable and practical. While prior cancer cell line alignment frameworks, such as Celligner,^50^ used bulk RNA-seq to compare primary tumors and cell lines, the scTumor Atlas enables direct single-cell comparisons to isolate malignant cell profiles. Finally, by linking atlas-derived transcriptional states to high-throughput cell line-derived dependencies, this work advances prior dependency mapping efforts that relied on bulk transcriptomic profiles,^35^ providing a direct bridge between primary cancer cells *in vivo* and cancer cell lines *in vitro*.

In contrast to many contemporary atlas efforts that emphasize the collection of large numbers of cells, often at the expense of transcriptional depth and interpretability, the scTumor Atlas adopts a fundamentally different strategy centered on high-confidence malignant state representation. Large pan-cancer atlases and integration benchmarking have demonstrated that the scRNA-seq data included are frequently sparse and dominated by low-count cells, a limitation that complicates downstream modeling and contributes to inconsistent findings across cancer types.^12,51^ Many approaches to address this limitation focus on aggressive batch correction, transfer learning from reference models, or excessive data scaling, which often emphasize maximal inclusion and preservation of rare cell populations.^21,52,53^ Our approach was explicitly designed to yield a compact yet biologically coherent reference that avoids the distortions introduced by low-depth profiles and outlier populations. Finally, while prior alignment frameworks evaluated bulk tumor to cell line concordance,^35^ mapping confidence decreases rapidly as nonmalignant cell fractions increase. The next incremental step to address this would intuitively be to map bulk cell line RNA-seq data to scRNA-seq in tumors, but this approach suffers considerably from bulk RNA-seq to scRNA-seq integration artifacts. By enabling direct scRNA-seq to scRNA-seq comparisons, the scTumor Atlas preserves biological structure at single-cell resolution and provides a more faithful foundation for evaluating model concordance and inferring functional dependencies.

This study has several limitations that should be acknowledged. Atlas composition is constrained by the availability and quality of public scRNA-seq datasets, resulting in uneven representation across cancer types. While cell number limitations partially address this issue, sample numbers remain skewed toward extensively published solid tumors, such as lung, pancreas, and prostate cancers. Our downsampling strategy was intentionally designed to prioritize inclusion of the most representative malignant cell states within each cancer type, enabling robust cross-cancer integration and comparison. As a result, this approach deemphasizes finer-grained phenotypic heterogeneity within individual tumor types, such as hormone receptor–defined states in breast cancer or basal and classical programs in pancreatic cancer. Detailed resolution of these intra-tumor and subtype-specific transcriptional programs, while biologically important, requires targeted sampling strategies and tumor-specific analytical frameworks and was therefore considered beyond the scope of this pan-cancer atlas. Dependency models are trained on expression profiles obtained by summing single-cell measurements within each cancer cell line. Although commonly termed “pseudobulking,” this approach preserves a gene-level representation that remains naturally compatible with scRNA-seq tumor data and avoids the integration challenges inherent to conventional bulk cell line RNA-seq.^54^ Finally, the broader applicability of dependency modeling is currently constrained by the limited number of cancer cell lines with available scRNA-seq data. Historically, single-cell profiling of cell lines was pursued primarily to assess intra-line heterogeneity, which is modest compared to primary tumors. Our findings suggest that continued single-cell profiling of cancer cell lines serves an additional and underappreciated role: improving the fidelity with which functional dependencies inferred from tumor scRNA-seq data may be modeled and interpreted.

In summary, this work introduces a large-scale, pan-cancer strategy for interpreting tumor biology across diverse malignancies. By integrating tumor identity, cell line concordance, and functional dependency prediction within a unified single-cell framework, the scTumor Atlas enables systematic and scalable translation of high-throughput functional genomics efforts. Future efforts will expand atlas breadth, incorporate spatial and multiomic data, and extend dependency modeling to patient-derived systems, supporting increasingly precise and context-aware therapeutic discovery.

## Methods

### Sample Annotation, Quality Control, and Merged Object Construction

Publicly available repositories containing raw single-cell RNA sequencing (scRNA-seq) count matrices were queried, including the Cancer Single Cell Expression Map (CancerSCEM),^7,47^ the Weizmann Curated Cancer Cell Atlas (WCCA),^5,8^ and the Gene Expression Omnibus (GEO),^55^ from which raw count matrices, barcode files, gene annotations, and associated metadata were downloaded (**Supplementary Table 1**). Datasets with alternative AnnData^56^ structures were standardized using a Scanpy workflow that included quality control, doublet detection, and dimensionality reduction.^57^ Cells were filtered based on total unique molecular identifier (UMI) count (≥1,000 and ≤5×10^6^), minimum cell count (≥200), and mitochondrial gene expression (≤10%). Doublets were identified and removed using Scrublet^19^ with an expected doublet rate of 5% (applied to samples containing at least 50 cells). Filtered datasets were then normalized and log-transformed prior to principal component analysis (PCA), neighborhood graph construction, UMAP embedding, and Leiden clustering. This pipeline was applied to the following datasets: ccRCC, PNET, HCC, DSRCT, SyS, OS, ALL, CRC, and CESC (**Supplementary Table 1**).

If malignant cell labels were available from the original dataset annotations, these labels were retained. In datasets lacking a clear or consistent malignant designation (PNET, HCC, DSRCT, CESC), cancer-specific classification rules were applied. First, cells were grouped into transcriptionally coherent communities using Leiden clustering at resolution 0.1. The distribution of pre-existing cell-type annotations, expression of lineage-associated markers (epithelial, fibroblast, smooth muscle, and leukocyte), and cancer-specific scoring criteria were jointly evaluated to classify clusters as malignant or non-malignant. For HCC, clusters with majority hepatocyte annotation were defined as malignant. For DSRCT, malignant cells were defined as the cluster level in which the expression of a DSRCT tumor marker set (*GALP, NGB, GAL, CCL25, GJB6, ITIH2*) exceeded the mean expression of immune/stromal markers (*C1QA, PTPRC, LUM, PECAM1, CD3E, JCHAIN*) by at least 0.5 (cluster-level score ≥0.5). For cervical squamous cell carcinoma and endocervical adenocarcinoma (CESC), malignant clusters were identified with a keratin-based score computed as the difference between mean *KRT8/KRT19* expression and mean immune/stromal marker expression, labeling clusters with score ≥0.5 as malignant (**Extended Data Fig. 1**).

Following the custom processing of samples noted above, all samples were reprocessed using a Scanpy workflow that restricted genes to a common reference gene list and excluded samples lacking valid sample identifiers or containing <200 cells. Cells with total UMI counts <5,000 or ≥5×10^6^ and mitochondrial gene expression >10% were also excluded. Doublet detection was reapplied as described above. Datasets were individually normalized and log transformed prior to PCA, neighbor graph construction, UMAP embedding (min_dist = 0.3), and Leiden clustering (resolution = 0.1). Broad cell types were confirmed by plotting the expression of the following markers: *PTPRC* (leukocytes), *KRT8* and *KRT18* (epithelial), *LUM* (fibroblast/stromal), *PECAM1* (endothelial), *PDGFRB* (pericytes/mesenchymal), and *TAGLN* (smooth muscle cells). Cancer types with multiple samples were batch-corrected and integrated using Harmony.^20^

Mahalanobis distance-based downsampling was performed at two levels: first at the sample level, and then at the cancer type level. First, at the sample level, malignant cells were subset and capped at 5,000 per sample via Mahalanobis distance-based downsampling for samples exceeding 5,000 malignant cells; samples with ≤5,000 malignant cells were retained without downsampling. Mahalanobis distances were calculated using the first 10 principal components derived from the top 2000 highly variable genes. Outliers were classified as Mahalanobis distances exceeding the 75th percentile of the chi-squared distribution (chi2_threshold = 0.75) and removed from the malignant population. The resulting subsample of at most 5,000 malignant cells for each sample was stored as an anndata object. Next, a second round of Mahalanobis distance-based downsampling was performed to downsample each cancer type to ≤5,000 representative cells. For each cancer type, the previously downsampled anndata objects were concatenated together and were subjected to log transformation, PCA, Harmony integration, and Mahalanobis distance-based downsampling using Harmony-adjusted principal components, following the same parameters as in sample-level downsampling. Once the two-step downsampling procedure was complete, the resulting cancer-type anndata objects were merged, and the concatenated anndata object was normalized and log transformed.

### ScVI and scANVI Tumor Atlas Construction

To construct a unified, batch-corrected, and semi-supervised integrated atlas, we applied an scVI/scANVI framework.^21,22^ Raw UMI counts were provided to scVI to learn an unsupervised probabilistic latent representation while correcting for sample-level batch effects, with samples containing fewer than 50 cells grouped into a shared batch category to ensure model stability. Building on the trained scVI model, scANVI was then used to incorporate cancer type labels in a semi-supervised manner, encouraging separation of known tumor classes while preserving the generative structure of the data. Training was performed for 200 epochs for scVI, followed by 100 epochs for scANVI. Models were trained on anndata objects with varying feature sets, including the top 2,000–5,000 highly variable genes as well as all genes. Of note, prior to selection of highly variable genes, the features were subset to those present in the bulk cancer cell line data to provide maximum feature overlap. After evaluation of the results from the different scANVI models, the model constructed from 3000 highly variable genes was chosen for downstream analysis because it offered the best balance of denoising the data while providing an adequate number of genes for alignment with cell lines and cancer dependency prediction. The scANVI latent space was used for downstream neighborhood graph construction and UMAP visualization. Visualization with UMAP, dot plots, and track plots was performed using Scanpy.

### Preranked GSEA

Rank-based pathway analysis was performed using gene set enrichment analysis (GSEA) in Python (GSEApy, https://github.com/zqfang/GSEApy) with the MSigDB Hallmark gene set collection (Hs_v2025.1).^58–60^ Genes were ranked by differential expression between groups using Scanpy’s Wilcoxon test, and preranked GSEA was performed with 5000 permutations. Pathway enrichment was quantified using normalized enrichment scores. Statistical significance was assessed by permutation testing, with multiple testing correction performed using the false discovery rate.

### Bulk RNA-Seq Data Download, Processing, and Analysis

TCGA bulk RNA-seq data from 8,246 tumors corresponding to cancer types represented in the atlas were analyzed (**Supplementary Table 7**). Bulk RNA-seq expression data for primary tumors (TCGA) and cancer cell lines (CCL) were obtained from the Celligner repository (https://github.com/broadinstitute/._ms) as transcript-per-million matrices.^50^ Gene names were standardized using the Human Genome Organization Genome Nomenclature Committee (HGNC) reference set, and analyses were restricted to shared protein-coding genes. TCGA and CCL datasets were converted into AnnData objects, and available sample annotations were incorporated from the Celligner metadata. Samples without valid annotations were excluded. For bulk tumor samples, 3,000 highly variable genes were selected, followed by scaling, dimension reduction, neighborhood graph construction, and UMAP embedding. Bulk cancer–specific gene sets were defined as the top 100 positively differentially expressed genes per cancer type using the scanpy.tl.rank_genes_groups() function (Wilcoxon method, log fold change >0, *p_adj_* <0.05).

Bulk CCL RNA-seq expression matrices and associated Celligner metadata were downloaded from the Celligner repository and filtered to HGNC-approved gene symbols by intersecting features with the HGNC complete set and removing duplicate entries. Disease/subtype annotations were mapped to atlas cancer-type abbreviations using curated reference tables. Samples lacking valid mappings excluded.

### Single Cell Cancer Cell Line (scCCL) Data Download, Processing and Analysis

ScRNA-seq CCL count matrices were downloaded as raw counts and converted to AnnData objects, with gene identifiers harmonized to gene symbols where necessary (**Supplementary Table 1**).^49,61^ Single-cell data were subjected to standard quality control, including removal of low-coverage cells (<2,000 counts) and high mitochondrial-content cells (>10%). Predicted doublets were identified using Scrublet, with an expected doublet rate of 5%.^19^ Preprocessed single-cell data were normalized and log-transformed prior to downstream analyses.

To quantify cross-modality concordance between bulk and single-cell CCL expression profiles, canonical correlation analysis (CCA) was performed on matched cell lines. Using decoupler,^62^ Single-cell RNA-seq data were aggregated into pseudobulk profiles by summing the raw counts per DepMap cell line, followed by normalization, log transformation, and scaling. Bulk CCL expression profiles were matched to the same DepMap identifiers and processed on a shared gene space using scaled log_2_(TPM) values. Both pseudobulk single-cell and bulk matrices were restricted to intersecting genes and matched cell lines, and CCA was performed with scikit-learn using five components.^63^ Pairwise similarity between modalities was assessed using cosine similarity computed between CCA embeddings for each matched cell line and visualized as a similarity heatmap (**Fig. 5C**). Two-dimensional CCA projections were additionally visualized by plotting the first two canonical components for both modalities, with matched bulk and pseudobulk points connected by line segments to illustrate per–cell line concordance across modalities (**Fig. 5D**). Cancer-type–specific gene sets were generated from scCCL data, retaining the top 100 positively enriched genes per scCCL type (p_adj_<0.05). These scCCL gene sets were used to demonstrate concordance with scTumor Atlas (**Fig. 5G**).

CCL scRNA-seq profiles were embedded into the tumor atlas latent space by further training the scTumor scANVI model on the loaded query scCCL anndata object for 200 epochs. Latent representations of CCL were overlaid onto the existing tumor atlas UMAP by first identifying the 15 nearest neighbor cells using Euclidean distance in scANVI latent space and then assigning each cell line a UMAP position based on a distance-weighted average of those tumor cells’ UMAP coordinates. The resulting cell line positions were summarized by computing embedding density across the UMAP using Scanpy’s embedding_density() function, producing a continuous map highlighting regions where cells were most concentrated. Tumor atlas cells were shown as low-opacity background points for reference (**Fig. 5H**). To compute normalized latent space distances between scCCL and scTumor Atlas primary tumors, tumor atlas cells were used to define a latent-space centroid by averaging scANVI embeddings, and each matched cell line was represented by its own centroid embedding. The Euclidean distance from each cell line centroid to the corresponding tumor centroid was computed (**Fig. 5K**).

### scTumor Atlas Model Validation in Breast Cancer

An independent scRNA-seq dataset of breast cancer cell lines^34^ was downloaded as raw count matrices and converted to AnnData objects. Gene identifiers provided as Ensembl IDs were mapped to HGNC gene symbols prior to analysis. Cells were filtered to remove low-quality (<2,000 total counts) and high mitochondrial-content cells (>10%). Doublets were removed using Scrublet, with an expected doublet rate of 5%.^19^ The anndata objects were then normalized and log transformed. Preprocessed breast cancer cell lines were embedded into the scTumor Atlas by further training the scANVI model for 20 epochs without supplying cancer-type labels. Latent representations were projected onto the existing scTumor Atlas UMAP as described above.

### DepMap CRISPR Screen

We adapted a previously described dependency mapping approach to our single-cell dataset.^35^ CRISPR–Cas9 gene dependency data were obtained from the Broad Institute DepMap Public 25Q3 release (https://depmap.org/portal).^17,37^ Gene-effect scores inferred using the Chronos model were used as ground-truth dependency estimates, with more negative values indicating reduced cellular fitness following gene knockout. Pseudobulked RNA-sequencing expression profiles for DepMap cancer cell lines were derived from matched single-cell CCL datasets by aggregating raw counts per cell line, followed by library-size normalization and log transformation. To ensure strong dependency predictions, only genes with at least 2 strong dependencies (Chronos score < −1 in ≥2 cell lines) and at least 2 independencies (Chronos score > −0.2 in ≥2 cell lines) were retained. Prior to model fitting, gene expression values were standardized across cell lines using z-score transformation (zero mean, unit variance) to place all genes on a comparable scale.

For each candidate dependency gene, an ElasticNet regression model was trained to predict Chronos gene-effect scores (predicted gene effect score, PGES) on each single cell in the scTumor Atlas using single-cell–derived CCL expression profiles as input features. Models were trained independently per gene using cross-validated ElasticNet regression (ElasticNetCV of scikit-learn^63^), with L1/L2 mixing parameters (l1_ratio ∈ {0.1, 0.5, 0.9}) and regularization strength selected via 10-fold cross-validation.^35,36^ Genes with Chronos gene-effect scores available in <40 cell lines were excluded from model training. Model performance was evaluated using the coefficient of determination (R²) on the training data. Only dependency models achieving R² ≥0.5 were retained for downstream analyses, yielding a high-confidence set of 529 predictive dependency genes. The DepMap DEMETER2 framework was used to model gene dependency by RNAi.^42^ Additional scRNA-seq datasets were incorporated by computing z-scored expression features prior to ElasticNet prediction and PGES estimation (e.g., RPLMS).

### Sample Preparation of RPLMS for Single-cell RNA Sequencing

Primary tumors were dissociated into single cells using the Miltenyi Human Tumor Dissociation Kit (#130-095-929) and stored at −80 °C until sequencing. ScRNA-seq libraries were created by Retro Biosciences with 10x Next GEM 5’ High Throughput workflow per protocol and then sequenced on an Illumina NovaSeq through Novogene, targeting at least 20,000 reads per cell.

### Single-cell Alignment, Preprocessing, and Annotation of RPLMS scRNA-seq Data

The RPLMS FASTQs were aligned using cellranger^64^ v7.1.0 to reference genome GRCh38-2020-A. The anndata object was filtered for cells with at least 2000 counts and less than 10% mitochondrial transcripts. Doublets were removed with Scrublet with an expected doublet rate of 5%. Normalization, log transformation, feature selection, PCA, neighborhood graph construction, Leiden clustering (resolution = 0.5) and UMAP (min_dist = 1) were performed using Scanpy. Clusters were manually annotated using canonical markers. One cluster that consisted of doublets was removed. The anndata was then filtered for tumor cells and a shared gene list prior to integration using scANVI.

### Single-Cell CRISPR Dependency Validation in RPLMS

RPLMS cells were integrated into the pretrained scTumor Atlas using scANVI in fully unlabeled query mode with further training for 50 epochs and visualized on the existing tumor atlas UMAP using the previously described latent-space nearest-neighbor projection approach. ElasticNet dependency models trained on DepMap Chronos CRISPR–Cas9 screens were applied to RPLMS single-cell expression profiles to generate predicted gene-effect scores. Predicted dependencies were combined with atlas-derived predictions, z-score normalized, and analyzed using Wilcoxon rank sum testing to identify RPLMS–selective dependencies.

## Data and code availability

RPLMS scRNA-seq data have been deposited on GEO with the accession number GSE319327. The scTumor scANVI model and post-scANVI anndata object are available at https://doi.org/10.5281/zenodo.18625030. All code from this paper is available at https://github.com/delittolab/scTumor and will be made publicly accessible upon the completion of peer review.

## Acknowledgements

D.D. was supported by funding from the Damon Runyon Cancer Research Foundation, the American Cancer Society, the National Institutes of Health (NIH) Loan Repayment Program, and the John and Marva Warnock Faculty Scholar Award. D.F.S., R.F.R.T., M.K. and C.E.G. were supported by the Division of General Surgery, Department of Surgery, School of Medicine, Stanford University. M.K. and J.P.A. were supported by the Transplant Tissue Engineering Center of Excellence at Stanford University. M.K. was also supported by the Advanced Residency Training at Stanford Program. R.F.R.T. was also supported by the NIH Loan Repayment Program.

M.T.L was supported by the National Institutes of Health (NIH), the U.S. Department of Defense, the Scleroderma Research Foundation, and the Wu Tsai Human Performance Alliance. Computational analyses were supported by the Stanford Research Computing Center at Stanford University and Retro Biosciences. The authors would like to acknowledge Chris Jeon for his work on the GPU partition of the Stanford Computational Genomics (SCG) Cluster. The authors would like to acknowledge Retro Biosciences for performing the 10x Genomics library preparation and sequencing of the RPLMS tumor sample. The results shown here are in part based upon data generated by the TCGA Research Network (https://www.cancer.gov/tcga) and DepMap (https://depmap.org/portal).

## Author contributions

Conceptualization, funding acquisition, resources, supervision, D.D.; Investigation, R.F.R.T., D.S.F., D.D.; Formal analysis, R.F.R.T., J.P.A., C.G., D.S.F, D.D.; Writing - original draft, R.F.R.T, J.P.A., D.S.F., D.D.; Writing - review & editing, all authors.

## Competing interests

The author(s) declare no competing interests.

---

**Extended Data Fig. 1.**
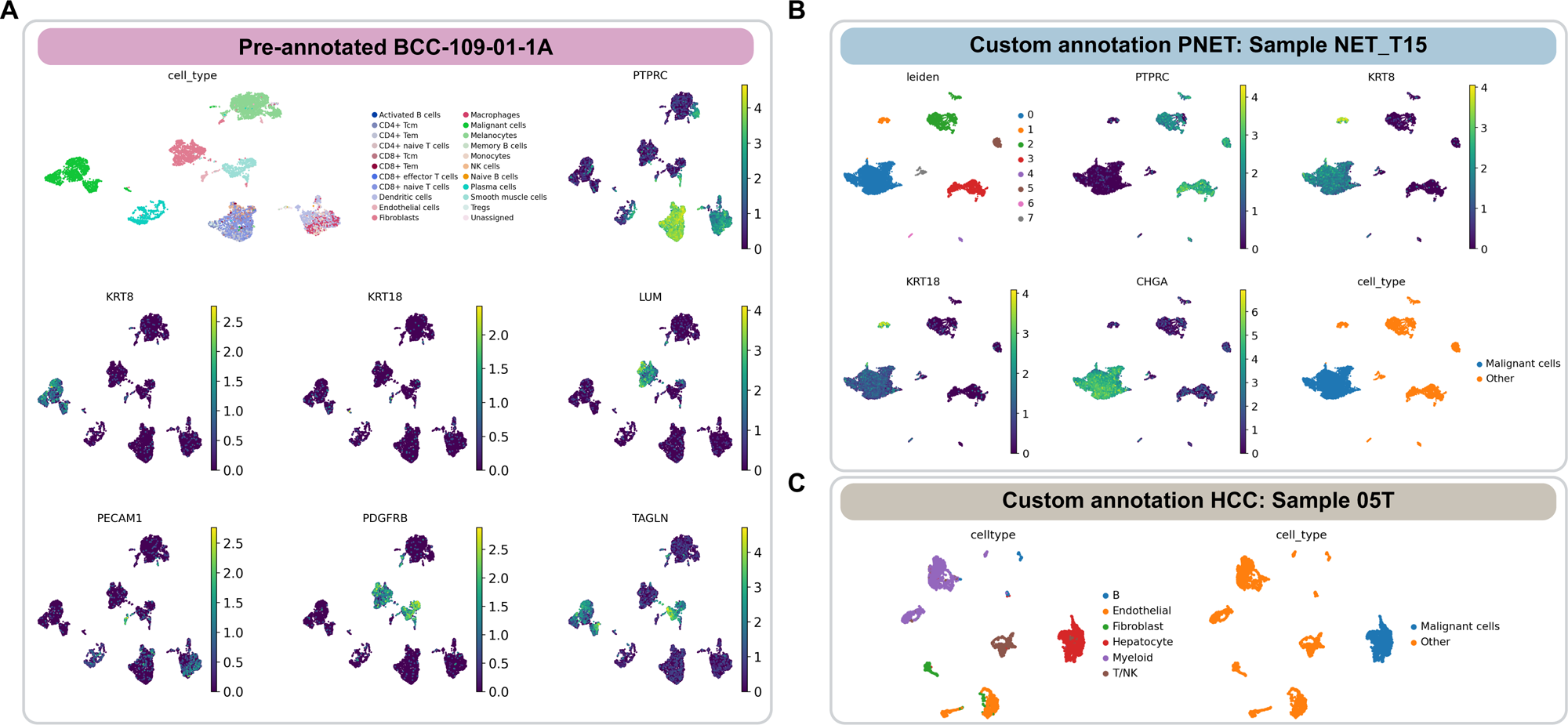
scTumor Atlas Annotation. Individual cancer type samples were grouped into transcriptionally coherent communities using Leiden clustering (resolution = 0.1). (**A**) Uniform Manifold Approximation and Projection (UMAP) of a representative pre-annotated basal cell carcinoma (BCC) dataset with original cell-type labels, with expression of canonical marker genes confirming expected tumor cell identity. (**B**) Custom annotation of a pancreatic neuroendocrine tumor (PNET) sample. Clusters were classified as malignant if their average *CHGA* expression exceeded 0.5. (**C**) Custom annotation of a hepatocellular carcinoma (HCC) sample. Clusters in which the dominant cell-type label was originally labeled “Hepatocyte” were reclassified as “Malignant,” while remaining clusters were relabeled as “Other.”

**Extended Data Fig. 2.**
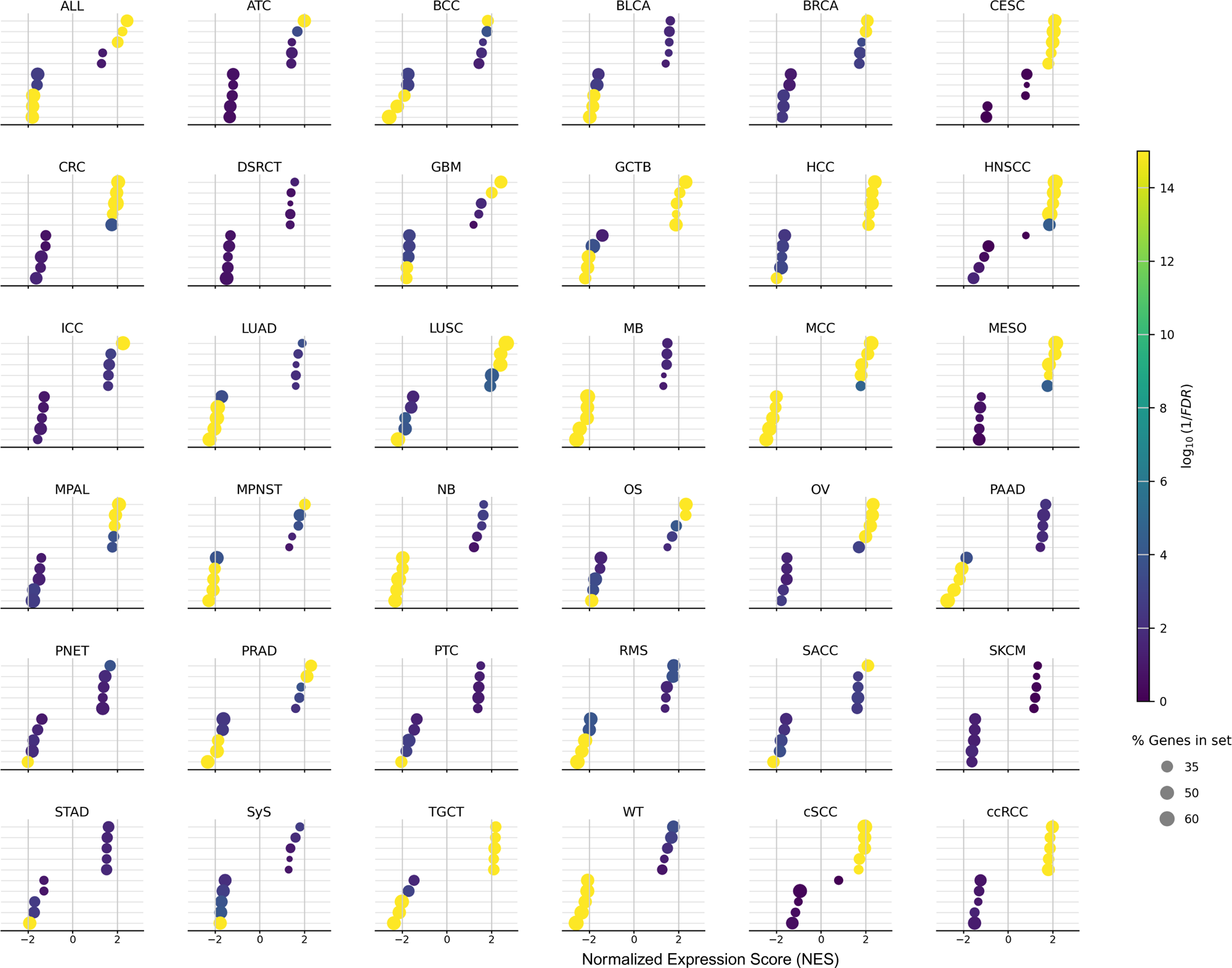
Top 5 Positively and Negatively Enriched GSEA Hallmark Pathways Across scTumor Atlas Cancer Types. Combined dot plot showing the top five positively and negatively enriched Hallmark gene sets for each cancer type in the scTumor Atlas, ranked by normalized enrichment score (NES). Dot color represents statistical significance (log₁₀(1/FDR)) while dot size indicates percent gene contribution to pathway (gene%) (**Supplementary Table 3**). Cancer type abbreviations are given in Fig. 1A.

**Extended Data Fig. 3.**
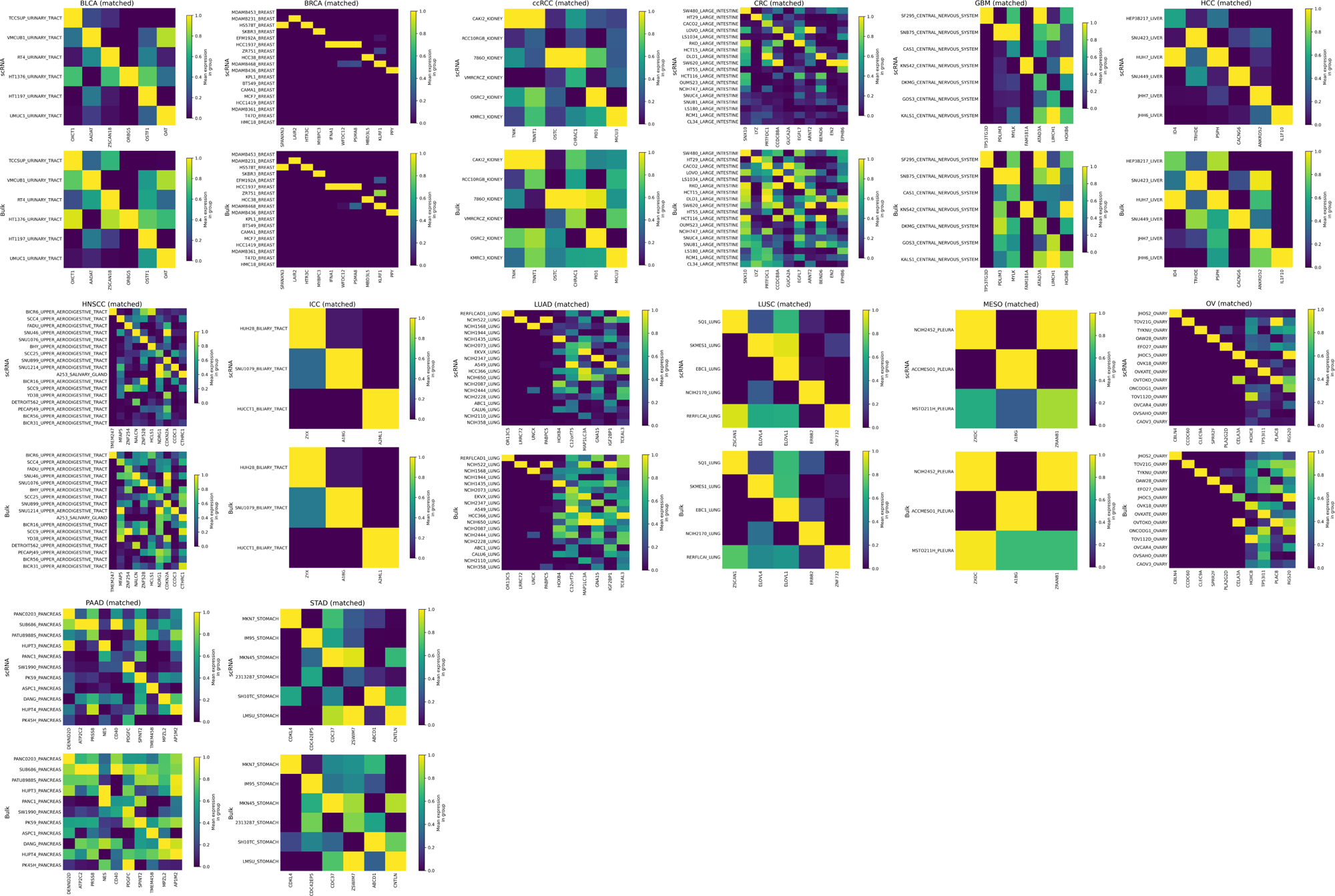
scRNA-seq and Bulk RNA-seq concordance in matched cancer cell lines. Stacked heatmaps illustrating the top 10 genes per cancer type ranked by Spearman correlation between pseudobulked single-cell RNA-seq and bulk RNA-seq expression across matched cancer cell lines. Pseudobulked single-cell expression (top) and bulk RNA-seq expression (bottom) are shown for each cancer type. Expression values are scaled per gene (0–1). Rows (cell lines) and columns (genes) were reordered to emphasize concordant expression patterns between modalities.

**Extended Data Fig. 4.**
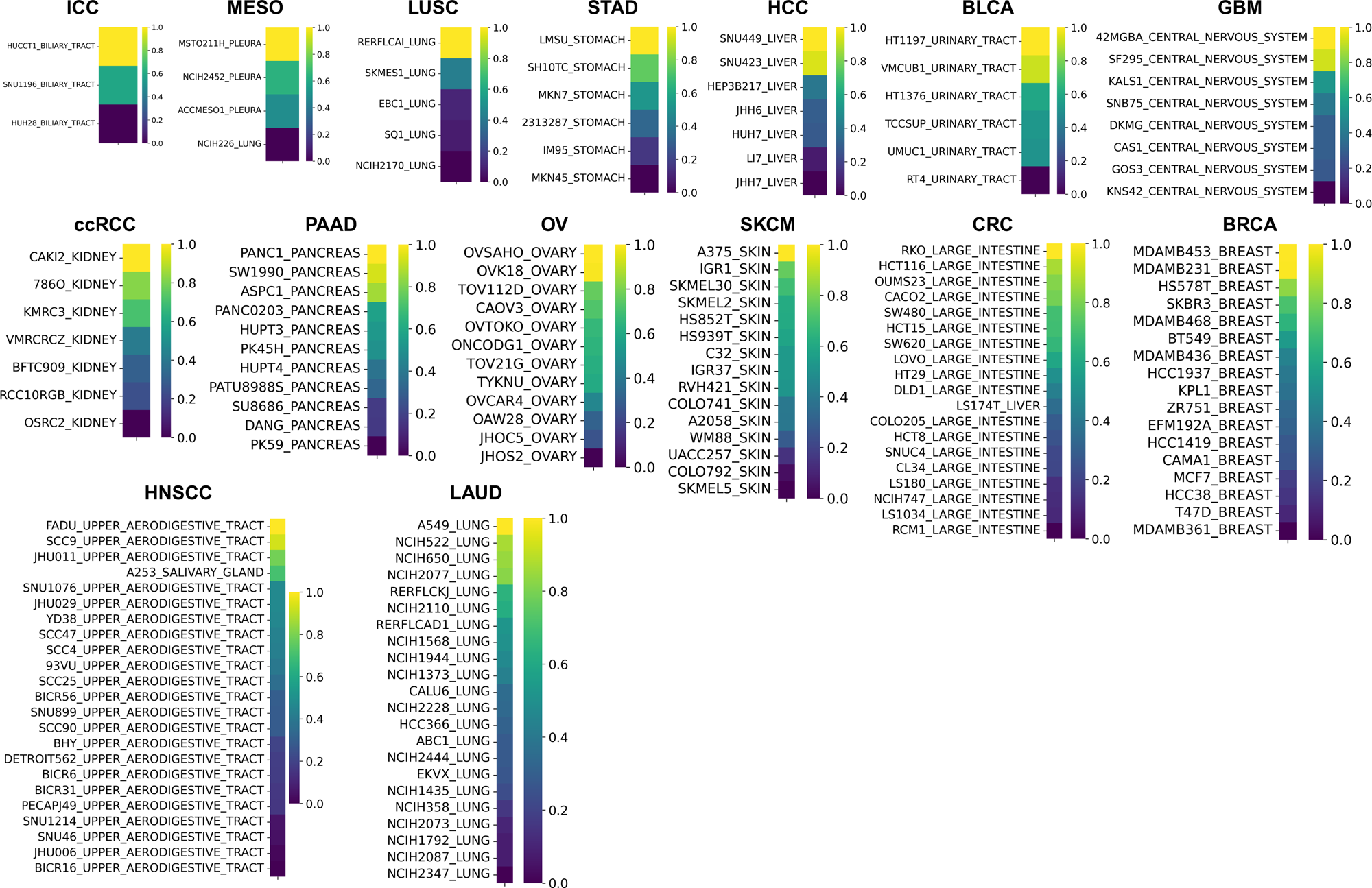
Normalized Latent Space Distances Between Single-Cell Tumor Atlas and Single-Cell Cancer Cell Lines. Heat map showing Euclidean distances between cancer cell line centroids and their corresponding scTumor Atlas (primary tumor) centroid. Distances were computed in the integrated latent space. Lower Euclidean distances (blue) indicate greater transcriptional similarity and higher fidelity of cell lines to their tumor counterparts, whereas higher distances (yellow) reflect increased divergence from primary tumor cell states. Only cancer types represented by more than two cell lines are shown.

**Extended Data Fig. 5.**
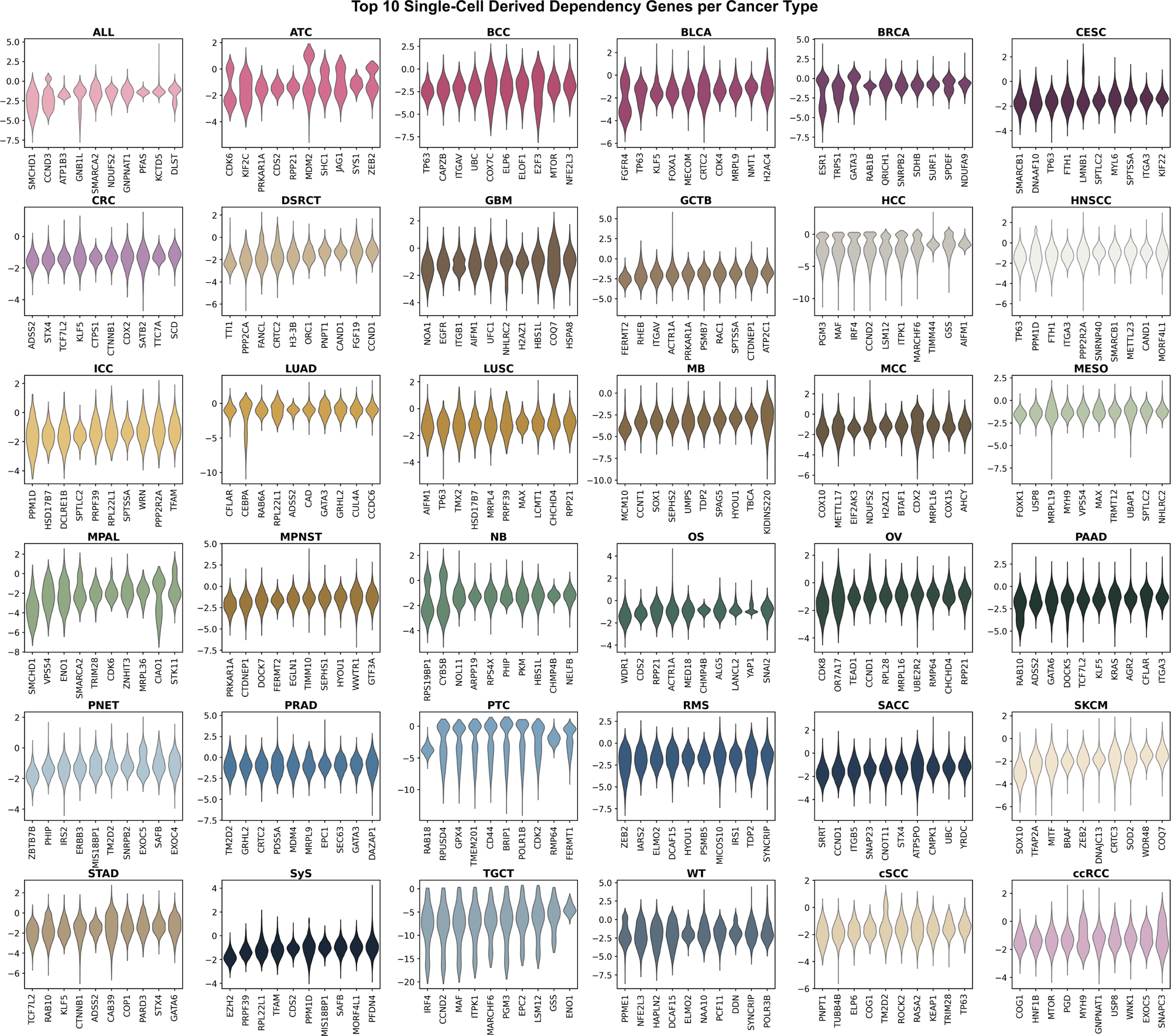
Single-Cell–Derived Gene Dependency Profiles Across Cancer Types. Violin plots of the top 10 single-cell–derived gene dependencies in Fig. 6 per scTumor Atlas cancer type. Genes were ranked by dependency z-score, with lower values indicating stronger dependency.

**Supplementary Table 1.** scTumor Atlas Sample Information

**Supplementary Table 2.** Top 50 Differentially Expressed Genes by ScTumor Atlas Cancer Type

**Supplementary Table 3.** Hallmark Pathway Enrichment Statistics for the Top 5 Positively and Negatively Enriched Pathways Across Cancer Types

**Supplementary Table 4.** Top 100 Ranked Genes in Tumor Bulk RNA-seq

**Supplementary Table 5.** Top 100 Ranked Genes in Cancer Cell Line scRNA-seq

**Supplementary Table 6.** A List of the ElasticNet-Model–Derived Gene Dependencies from scTumor Atlas

**Supplementary Table 7.** TCGA Bulk Tumor and Single-Cell Sequencing Sample Information

## Notes

https://github.com/delittolab/scTumor

### Competing Interest Statement

The authors have declared no competing interest.

### Summary of Updates

One author's name was misspelled; this was corrected.

