## Supplementary Table 1 for "A Pan-Cancer Single-Cell Atlas to Evaluate Tumor Identity, Cell Line Concordance, and Dependency Mapping"

**Supplementary Table 1.** Sample Sequencing Information

| *Sample* | *Reference* | *Database* | *Data Source* | *Accession Number* | *Datasets* |
|---|---|---|---|---|---|
| ***Bulk RNA-seq*** | | | | | |
| *Primary Tumors (Fig. 4)* | | TCGA | | Refer to Supplementary Table 7 | |
| *Cancer Cell Lines (Fig. 5)* | | TCGA | | Refer to Supplementary Table 7 | |
| ***scRNA-seq - Cancer Cell Line*** | | | | | |
| *Cancer Cell Lines (Fig. 5 & Fig. 6)* | Qionghua Zhu et al., 2023[1]<br>Kinker et al., 2020[2] | N/A | CNSA<br>GSA<br>GEO | CNP0004330<br>PRJCA021248<br>GSE157220 | |
| ***scRNA-seq – Primary Tumors*** | | | | | |
| *Acute Lymphoblastic Lymphoma (ALL)* | Caron et al., 2020[3] | Weizmann CCA | GEO | GSE132509 | ETV6.RUNX1.1-1.4<br>HHD.1 & HHD.2<br>PBMMC.2 & PBMMC.3<br>PRE-T.1& PRE-T.2 |
| *Anaplastic Thyroid Cancer (ATC)* | Gao et al., 2021[4] | CancerSCEM | GEO | GSE148673 | ATC-026-(01-05)-1A |
| *Basal Cell Carcinoma (BCC)* | Yost et al., 2019[5]<br>Ganier et al., 2024[6] | CancerSCEM | GEO<br>EBI | GSE123813<br>E-MTAB-13085 | BCC-018-(01-05)-1A<br>BCC-109-(01-08)-1A |
| *Bladder Cancer (BLCA)* | ai et al.,2021[7]<br>Minoli et al., 2023[8] | CancerSCEM | GEO | GSE135337<br>GSE217956 | BLCA-031-(01-07)-1A<br>BLCA-115-(01-03)-1A |
| *Breast Cancer (BRCA)* | Qian et al.,2020[9]<br>Mao et al.,2022[10]<br>Cords et al.,2023[11]<br>Liu et al.,2023[12]<br>Song et al.,2023[13] | CancerSCEM | EBI<br>NGDC<br>GEO | E-MTAB-8107<br>PRJCA008495<br>E-MTAB-10607<br>GSE225600<br>GSE234832 | BRCA-030-(01-18)-1A<br>BRCA-071-(03,06,09,12)-1A<br>BRCA-107-(01-14)-1A<br>BRCA-128-(01-08)-1A<br>BRCA-134-(01-03)-1A |
| | Gambardella et al., 2022[14] | N/A | GEO | GSE173634 | AU565, BT20, BT474, BT483, CAL51, CAL851, DU4475, EVSAT, HCC(70, 1143, 1187, 1500, 1954), HDQP1, JIMT1, MDAMB415, MX1 |
| *Cervical squamous cell carcinoma (CESC)* | Li et al., 2022[15] | ArrayExpress | EBI | E-MTAB-11948 | Sample1-3 |
| *Clear Cell Renal Carcinoma (ccRCC)* | Bi et al., 2021[16]<br>Krishna et al., 2021[17] | Weizmann CCA | dbGaP<br>SRA | phs002065.v1.p1<br>PRJNA705464 | P55, 76, 90, 912, 913, 915, 916<br>UT1 & 2T1-4 |
| | Davidson et al., 2023[18]<br>Alchachin 2022[19]<br>Mei et al., 2024[20] | Cancer SCEM | GEO | GSE210038<br>GSE178481<br>GSE202813 | ccRCC-060-(01-07)-1A<br>ccRCC-061-(01-15)-1A<br>ccRCC-118-(01-08)-1A |
| *Colorectal Cancer (CRC)* | Qian et al., 2029[21]<br>Becker et al.,2022[22]<br>Fleischer et al.,2023[23]<br>Bala et al.,2023[24]<br>Yang et al., 2023[25] | CancerSCEM | EBI<br>GEO | E-MTAB-8107<br>GSE201348<br>GSE222300<br>E-MTAB-12022<br>GSE201348<br>GSE224679<br>GSE232525 | CRC-030-(01-14)-1A<br>CRC-067-(01-04)-1A<br>CRC-073-(01-02)-1A<br>CRC-086-(01-04)-1A<br>CRC-101-(01-05)-1A<br>CRC-119-(01)-1A<br>CRC-133-(01-02)-1A |
| | Lee et al., 2020[26] | Weizmann CCA | EGA | EGAS00001003779<br>EGAS00001003769 | SMC01,02,04,07,08,09,11,14,15,16,18,20,21,23,25 |
| *Cutaneous Squamous Cell Carcinoma (cSCC)* | Zou et al., 2023[27] | CancerSCEM | GEO | GSE193304 | cSCC-099-(01-03)-1A |

| | | | | | |
|---|---|---|---|---|---|
| *Desmoplastic Small Round Cell Tumor (DSRCT)* | Henon et al., 2024[28] | Geo | GEO | GSE263523 | GR2 1-4<br>IC 1-3<br>GR7 1 & 2 |
| *Giant Cell Tumor of Bone (GCTB)* | Yang et al., 2022[29]<br>Van Ijzendoorn et al. 2022[30] | CancerSCEM | GEO | GSE212341<br>GSE210750 | GCTB-045-(01-02)-1A<br>GCTB-052-(01-03)-1A |
| *Glioblastoma (GBM)* | Wang et al., 2020[31]<br>Al-Dalahmah et al., 2023[32]<br>Park et al., 2022[33] | CancerSCEM | EBI<br>GEO | GSE139448<br>GSE224149<br>E-MTAB-12528<br>GSE189650 | GBM-010-(01-03)-1A<br>GBM-085-(01-18)-1A<br>GBM-108-(01-04)-1A<br>GBM-117-(01-02)-1A |
| *Head and Neck Squamous Cell Carcinoma (HNSCC)* | Lin et al., 2022[34]<br>Chu et al., 2023[35]<br>Chung et al., 2023[36] | CancerSCEM | GEO | GSE213047<br>GSE243359<br>GSE173468<br>GSE185965 | HNSCC-043-(01-03)-1A<br>HNSCC-087-(01-02)-1A<br>HNSCC-121-(01-15)-1A<br>HNSCC-127-(01-03)-1A |
| *Hepatocellular Carcinoma (HCC)* | Zhu et al., 2023[37]<br>Lu et al., 2022[38] | CancerSCEM | GEO | GSE202642<br>GSE155481<br>GSE149614 | HCC-059-(01-06)-1A<br>HCC-123-01-1A |
| *Intrahepatic Cholangiocarcinoma (ICC)* | Shi et al., 2022[39] | Cancer SCEM | GEO | GSE201425 | iCCA-066-01-1A<br>iCCA-066-(03-06)-1A |
| *Lung Adenocarcinoma (LUAD)* | Laughney et al., 2020[40]<br>Lambrechts et al., 2018[41]<br>Song et al., 2023[13] | CancerSCEM | GEO<br>EBI<br>SRA | GSE123904<br>E-MTAB-6149<br>PRJNA1055415<br>GSE234832 | LUAD-003-(01-13)-1A<br>LUAD-004-(01-03)-1A<br>LUAD-004-(06-08)-1A<br>LUAD-096-(01-56)-1A<br>LAUD-134-01-1A |
| *Lung Squamous Cell Carcinoma (LUSC)* | Lambrechts et al., 2018[41] | CancerSCEM | EBI<br>SRA | E-MTAB-6149<br>PRJNA976462 | LUSC-005-(01-06)-1A<br>LUSC-092-(01-26)-1A |
| *Malignant Peripheral Nerve Sheath Tumor (MPNST)* | Wu et al.,2022[42] | CancerSCEM<br>Geo | GEO | GSE179033 | MPNST-114-(01-04)-1A |
| *Medulloblastoma (MB)* | Funke et al., 2023[43]<br>Gold et al., 2024[44] | CancerSCEM | GEO | GSE212559<br>GSE214469 | MB-078-(01-06)-1A<br>MB-081-(01-13)-1A |
| *Merkel Cell Carcinoma (MCC)* | Paulson et al., 2018[45]<br>Das et al., 2023[46] | CancerScem<br>Geo<br>SRA | GEO<br>SRA | GSE117988<br>GSE118056<br>PRJNA1019891<br>GSE226438 | MCC-001-(01-02)-1A<br>MCC-002-(01-02)-1A<br>MCC-084-(01-10)-1A<br>MCC-122-(01-11)-1A |
| *Mesothelioma (MESO)* | Knelson et al., 2022[47] | CancerSCEM<br>GEO | GEO | GSE201925 | MESO-065-(01-03)-1A |
| *Mixed Phenotype Acute Leukemia (MPAL)* | Granja et al.,2019[48] | Geo | GEO | GSE139369 | MPAL-022-(01-10)-1A |
| *Neuroblastoma (NB)* | Dong et al., 2020[49] | CancerSCEM | GEO | GSE137804 | NB-033-(1-16)-1 |
| *Osteosarcoma (OS)* | Zhou et al., 2020[50] | CancerSCEM | GEO | GSE152048. | BC 2,3,5,6,10,11,16,17<br>BC 20-22 |
| *Ovarian Carcinoma (OV)* | Nelson et al., 2020[51]<br>Qian et al., 2020[21] | CancerSCEM | EBI | E-MTAB-8559<br>E-MTAB-8107 | OV-020-(01-04)-1A<br>OV-030-(01-07)-1A |
| *Pancreatic Adenocarcinoma (PAAD)* | Peng et al., 2019[52]<br>Chen et al., 2023[53]<br>Chen et al., 2023[54]<br>Schalck et al., 2022[55]<br>Shiau et al., 2022[56]<br>Storrs et al., 2023[57]<br>Zhang et al., 2023[58] | CancerSCEM | NGDC<br>(GSA)<br>SRA<br>GEO | PRJCA001063<br>PRJNA879876<br>GSE212966<br>GSE211644<br>GSE202051<br>GSE242230<br>PRJCA016878 | PDAC-027-(01-24)-1A<br>PDAC-044-(01-06)-1A<br>PDAC-046-(01-06)-1A<br>PDAC-051-(01-05)-1A<br>PDAC-064-(01-60)-1A<br>PDAC-091-(01-22)-1A<br>PDAC-106-(01-07)-1A |

| | | | | | |
|---|---|---|---|---|---|
| | Liu et al., 2023[59]<br>Oh et al., 2023[60] | | | GSE226762<br>GSE231535 | PDAC-129-01-1A<br>PDAC-132-(01-02)-1A |
| *Pancreatic Neuroendocrine Tumor (PNET)* | Ye et al., 2024[61] | N/A | GEO | GSE256136 | T1-15 |
| *Papillary Thyroid Carcinoma (PTC)* | Wang et al., 2022[62] | CancerSCEM | GEO | GSE191288 | PTC-069-01-1A |
| *Prostate Adenocarcinoma (PRAD)* | Chan et al.,2022[63]<br>Kfoury et al.,2021[64]<br>Hirz et al.,2023[65]<br>Pakulai et al.,2024[66] | CancerSCEM | GEO | GSE210358<br>GSE143791<br>GSE181294<br>GSE244267 | PRAD-053-(01-13)-1A<br>PRAD-063-(01-17)-1A<br>PRAD-072-(01-19)-1A<br>PRAD-082-(01-29)-1A |
| *Rhabdomyosarcoma (RMS)* | Patel et al.,2022[67] | CancerSCEM | GEO | GSE174376 | RMS-113-01- (1-20)-1A |
| *Salivary Adenoid Cystic Carcinoma (SACC)* | Zhou et al.,2023[68] | CancerSCEM | GEO | GSE217084 | SACC-103-(01-04)-1A |
| *Skin Cutaneous Melanoma (SKCM)* | Wu et al., 2024[69] | CancerSCEM | SRA | PRJNA996416 | SKCM-095-(01-03)-1A |
| *Stomach Adenocarcinoma (STAD)* | Alcindor et al., 2025[70] | CancerSCEM | SRA | PRJNA1052389 | GEA-093-(01-20)-1A |
| *Synovial Sarcoma (SyS)* | Jerby-Arnon et al., 2021[71] | Weizmann CCA | DUOS | DUOS-000123 | SyS1-2<br>SyS5<br>SyS7<br>SyS10-14<br>SyS16 |
| *Tenosynovial Giant Cell Tumor (TGCT)* | van Ijzendoorn et al., 2022[30] | CancerSCEM | GEO | GSE210750 | TGCT-052-(01-05)-1A |
| *Wilms Tumor (WT)* | Petrosyan et al., 2023[72] | CancerSCEM | GEO | GSE200256<br>GSE175698 | WT-070-(1-40)-1A<br>WT-124-01-1A |

*ArrayExpress (EBI), Cancer Single-cell Expression Map (CancerSCEM), China National GeneBank DataBase (CNGBdb) Sequence Archive (CNSA), Data Use Oversight System - Broad Institute (DUOS), European Genome-phenome Archive (EGA), Genome Sequence Archive (GSA), National Center for Biotechnology Information Gene Expression Omnibus (GEO), NIH database of Genotypes and Phenotypes (dbGaP), National Genomics Data Center (NGDC), National Center for Biotechnology Information Sequence Read Archive (SRA), Weizmann Curated Cancer Cell Atlas (Weizmann CCA)
